# Dendritic cell-derived lncRNA in patients with acute coronary syndrome

**DOI:** 10.1101/2023.12.05.570310

**Authors:** Zhenglong Wang, KE Changhao, Yuheng Chen, Yongchao Zhao, Yuanjie He, Xiao Liang, Qingxian Tu, Min Xu, Fujia Guo, Junbo Ge, Bei Shi

## Abstract

**Background:** Long non-coding RNA (lncRNA) and dendritic cells (DC) play an important role in the occurrence and development of acute coronary syndrome (ACS). However, the role and mechanism of DC-derived lncRNA in patients with ACS still remains understudied.

**Objective:** To investigate the expression difference and role of lncRNA in monocyte-derived DC(moDC) in patients with acute coronary syndrome(ACS).

**Methods and Results:** The subjects were divided into 4 groups: (1) healthy control group (CON), (2) unstable angina group (UA), (3) non-ST elevation myocardial infarction group (NSEMI or NST) and (4) ST elevation myocardial infarction group (STEMI or ST). Peripheral blood was extracted and isolated to obtain mononuclear cells (PBMC). After induction into moDC, morphology was observed by electron microscope and identified by flow cytometry (FCM) (CD80, CD86, CD11c, CD14, HLA-DR), and morphological and functional differences were compared. The differentially expressed lncRNA were screened by gene sequencing, and the biological functions of which were predicted by GO and KEGG and the correlation between candidate lncRNA and target genes was analyzed. The expression levels of markers, signaling pathways, inflammatory factors and target gene were performed by FCM, RT-PCR, WB and ELISA after the candidate lncRNA were over-expressed or silenced. Electron microscope showed that the cells were suspended cells of dendritic pseudopodia. FCM showed that the expression levels of DC markers CD11c (99.02±0.12%) and HLA-DR (99.32±0.08%) were high, while the expression levels of CD80 (24.27±0.99%) and CD86 (32.38±0.59%) were low. The number of differentially expressed lncRNA screened by gene sequencing was as follows: CON vs UA, 3; CON vs. NST, 49; CON vs ST, 35; UA vs. NST, 115; UA vs ST, 113; ST vs NST, 4. The results of FCM, RT-PCR, WB and ELISA showed that the expression levels of markers (CD80, CD86, HLA-DR), signaling pathways (p-PI3K, p-AKT), inflammatory factors (IL-6 and IL-12p70) and target gene (CCL14) were increased after over-expression of CCL15-CCL14; On the contrary, its expression level was decreased.

**Conclusions:** There are differences in moDC morphology and function and lncRNA expression in different types of ACS patients, and the differentially expressed lncRNA (CCL15-CCL14) regulates the function of moDC.

---

Acute coronary syndrome (ACS) is still a global problem dangerous to human health and life. At present, more studies on the pathogenesis of ACS focus on the local mechanisms of plaque characteristics and environmental factors^1–3^. However, Ounzain S et al hold that heart disease is the result of dysregulation of gene regulatory networks^4^, and hence, the regulatory factors of gene expression (rather than local and environmental factors) that lead to the pathogenesis and progression of ACS plaques are more of a concern^5^.

Long non-coding RNA(lncRNA) has previously been regarded as the “dark matter” and “noise” of genome transcription because it does not have biological functions^6^. However, many studies have confirmed that lncRNA is involved in the occurrence and process of ACS^7–9^. Meanwhile, related research hints that immunity plays an important role in atherosclerosis and ACS^10,11^. Based on the above, we speculated that the lncRNA of monocyte derived dendritic cells (moDC) were related to the occurrence and progression of ACS.

This study aimed to determine the role and mechanism of DC-derived lncRNA in patients with ACS, so as to provide a new perspective and reference for further exploring the pathogenesis of ACS.

## Methods

### Study patient

In present study, patients with acute coronary syndrome (ACS) were divided into four groups: (1) unstable angina (UA); (2) non-ST-segment elevation myocardial infarction (NSTEMI or NST); (3) ST-segment elevation myocardial infarction (STEMI or ST) and normal(control, CON). The protocol has been approved by the local ethics committee, and all patients signed an informed consent.

### moDC culture and identification

The peripheral blood(20ml) was obtained, and mononuclear cell (PBMC) was isolated by Ficoll density gradient centrifugation, and collected using an immunomagnetic bead isolation column(MACS) containing CD14 antibodies (Miltenly, Germany). 20ng/ml RATEGM-CSF (PeproTech, USA) and 20ng/ml rhIL-4rhIL-4 (PeproTech, USA) were added and placed in an incubator at 37℃ to induce PBMC into monocyte derived dendritic cells (moDC).

The identification of moDC morphology was performed by inverted phase contrast microscopy (OLYMPU, Germany), scanning electron microscopy (HITACHI, Japan) and transmission electron microscopy (HITACHI, Japan). The expression levels of moDC markers CD80(BioLegend, USA), CD86(BioLegend, USA) and HLA-DR (BioLegend, USA) were detected by flow cytometry (FCM).

### moDC morphological and functional differences

The morphological differences of moDC among different types of ACS patients were observed and compared by scanning electron microscopy. The expression levels of moDC markers CD80(BioLegend, USA), HLA-DR(BioLegend, USA) and ovalbumin (OVA) were detected by FCM to investigate the ability of antigen presentation and phagocytosis, and the expression levels of IL-6, IL10 and IL12p70 were detected by ELISA (Beijing Solarbio, China) to detect the ability of secretion.

### Differences in lncRNA expression

Illuminahiseq 3000 platform (RiboBio Co., Ltd., China) was used for sequencing to investigate the difference in lncRNA expression of moDC. The differentially expressed lncRNA were screened by significant level (Q value<0.05) and llog2(fold change)>1. The accuracy of differentially expressed lncRNA was verified by RT-PCR, and the biological functions of which were predicted by GO analysis and KEGG analysis. The pearson correlation coefficient was used to analyze the correlation between the expression level of candidate lncRNA and moDC markers (CD80, CD86, HLA-DR) as well as the expression levels of WBC, CTNT-hs, and CK-MB.

### Role and mechanism of candidate lncRNA

The functional changes of moDC, such as antigen presentation and cytokine secretion were performed by FCM and RT-PCR, and the phosphorylation level of moDC PI3K-AKT protein pathway was detected by western blot(WB) after over-expression or silence of candidate lncRNA in moDC. The sub-localization of candidate lncRNA in moDC was explored by fluorescence in situ hybridization, and The regulatory effect on the target gene was performed by FCM, RT-PCR and WB.

### Statistical analysis

SPSS29.0 statistical software was used for statistical analysis, Mann-Whitney test was used to compare continuous variables between the two groups, Χ2 test was used to compare qualitative data. Holm-Sidak method was used for comparison among multiple groups and one-way analysis of variance was performed. Linear regression and Spearman rank correlation were used to evaluate the correlation between lncRNA level and continuous variables, and logistic regression was used to evaluate the correlation between lncRNA level and binary classification variables. P<0.05 was defined as statistically significant difference^12^.

## Result

### Characteristics of the Study Population

Characteristics of the present study population (Table 1).

**Table 1.** Study Population Characteristics.

| variable | Control(CON)<br>(n=8) | STEMI(ST)<br>( n = 9 ) | NSTEMI(NST)<br>( n = 8 ) | UA<br>( n = 6 ) |
| --- | --- | --- | --- | --- |
| Age, y | 45.67±0.58 | 63.33±19.55 | 57.33±17.21 | 66.67±5.69 |
| SBP, mmHg | 117±6.25 | 142.33±15.63 | 146.67±30.55 | 127.33±23.59 |
| DBP, mmHg | 76.33±3.7C | 78.33±7.64 | 84.33±21.22 | 81.33±5.69 |
| HR,beats/min | 77±9.17 | 77±8.54 | 73.33±15.50 | 84±13.53 |
| WBC, ×10<sup>9</sup> /L | 8.45±4.40 | 9.62±3.47 | 9.78±3.74 | 8.7±1.58 |
| TC, mmol/L | 5.94±0.61 | 4.44±1.06* | 4.63±0.47* | 3.75±0.55* |
| TG, mmol/L | 2.68±1.16 | 2.17±0.94 | 3.42±2.10 | 1.22±0.32 |
| LDL-C, mmol/L | 3.60±0.21 | 2.83±0.78* | 2.77±0.23* | 2.35±0.58* |
| HDL-C, mmol/L | 1.39±0.31 | 1.04±0.10* | 1.03±0.15* | 1.09±0.29* |
| CK, U/L | 55.67±2.52 | 112.33±29.26 | 623.67±478.73 | 51.67±23.80 |
| CK-MB, U/L | 19.67±1436 | 21±1.0 | 44±3027 | 11.67±1.53 |
| TNT-hs, ng/L | 0.12±0.03 | 2799.95±4797.84 | 927.75±782.42 | 1.21±186 |
| Lipoprotein a, nmg/L | 108.02±59.56 | 212.41±283.52 | 71.2±74.51 | 88.27±67.47 |
| BNP, pg/ml | 51.11±21.58 | 2432.44±2209.07 | 623.03±656.14 | 61.91±5.47 |

Note: SBP, systolic blood pressure;DBP, diastolic blood pressure; WBC,White blood cells; HR,heart rate; Control, healthy volunteer; STEMI, ST-segment elevation myocardial infarction; NSTEMI, non-ST-segment elevation myocardial infarction; \*p<0.05, compared with CON.

### Identification of moDC

Inverted phase contrast microscopy showed that on the second day of culture, moDC grew in a semi-suspended and semi-adherent form (Figure 1A). On the 4th day of culture, moDC grew like suspension, with irregular shape and convex surface (Figure 1B). On the 6th day of culture, moDC showed duckweed aggregation, large volume, obvious dendritic protrusions and a large number of cell colonies (Figure 1C, D). Scanning electron microscopy (SEM) showed that moDC was a typical DC characteristics: elliptical shape, rough surface and laminated folds (Figure 2). Transmission electron microscopy showed that moDC was irregular in shape, with many protrudes of varying length and thickness at the edges, a large number of endoplasmic reticulum and mitochondria and a few lysosomes in the cytoplasm, and an irregular nucleus (Figure 3). The results of FCM showed that the levels of the markers CD11c (99.02±0.12%) and HLA-DR (99.32±0.08%) were high, and CD80 (24.27±0.99%) and CD86 (32.38±0.59%) were low (Figure 4).

**Figure 1.**
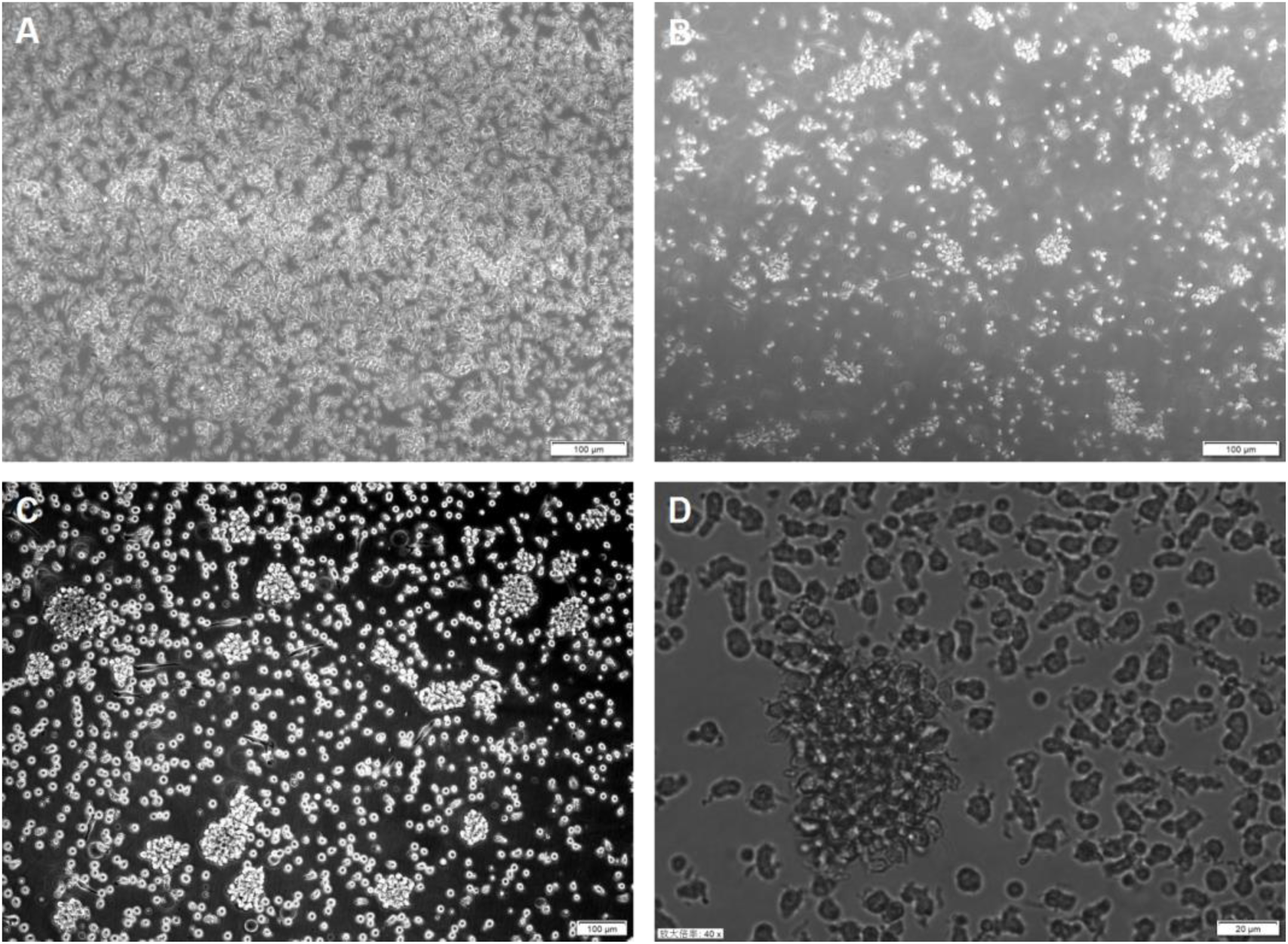
The moDC Morphology under Inverted Microscope. Note: A: Day 2 after MACS culture; B: Day 4 after MACS culture; C: Day 6 after MACS culture; A, B, C: scale =100μm; D: scale =20μm.

**Figure 2.**
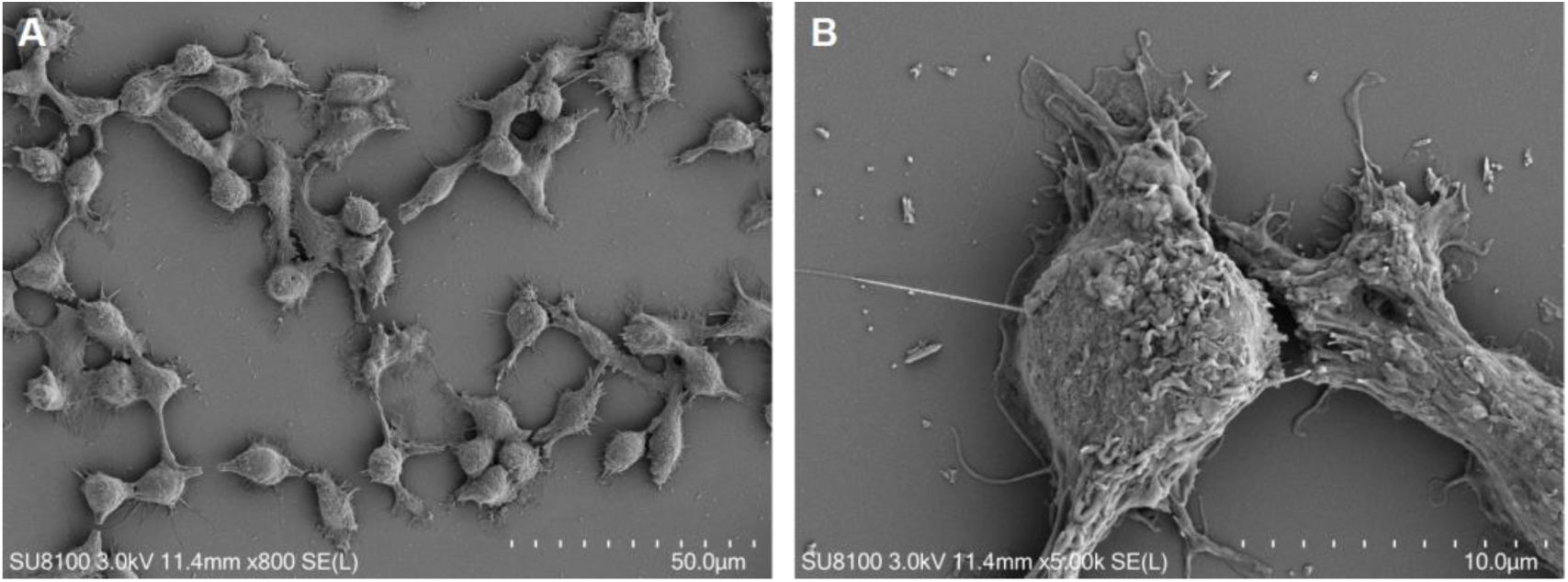
The moDC Morphology under Scanning Electron Microscope. Note: A: scale=50μm; B: scale=10μm.

**Figure 3.**
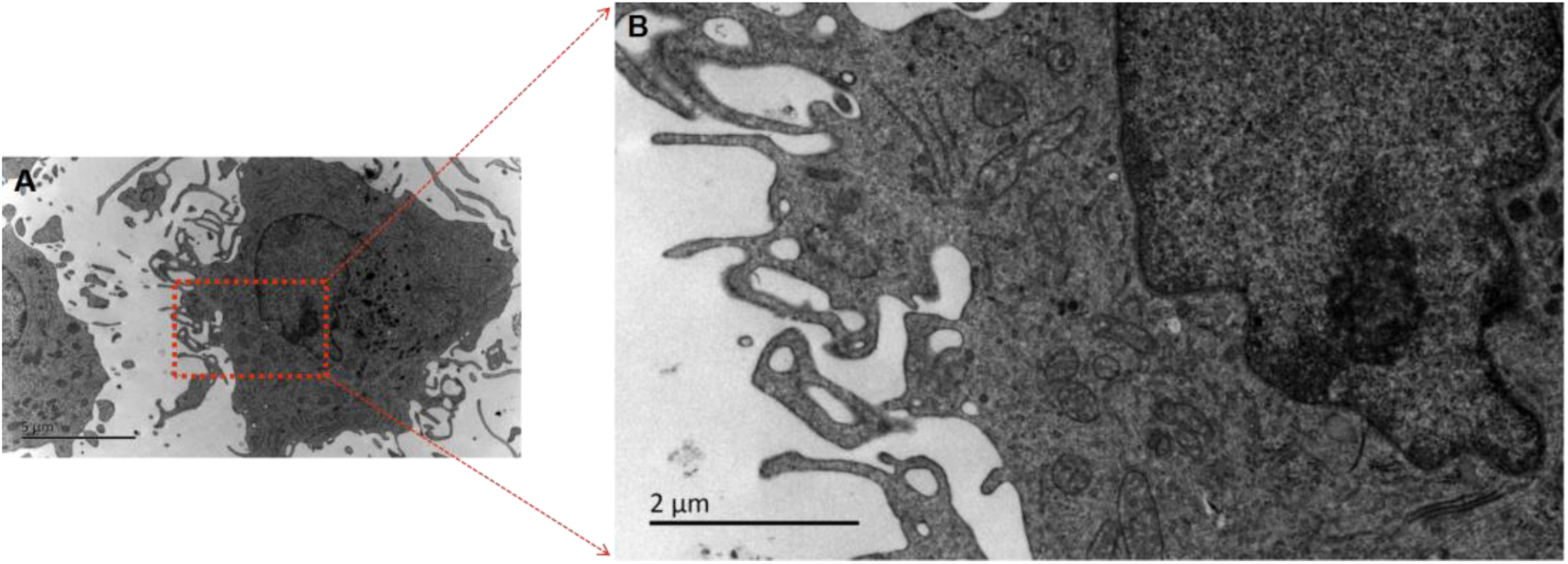
The moDC Morphology under Transmission Electron Microscope. Note: A: scale=5um; B: scale=2um.

**Figure 4.**
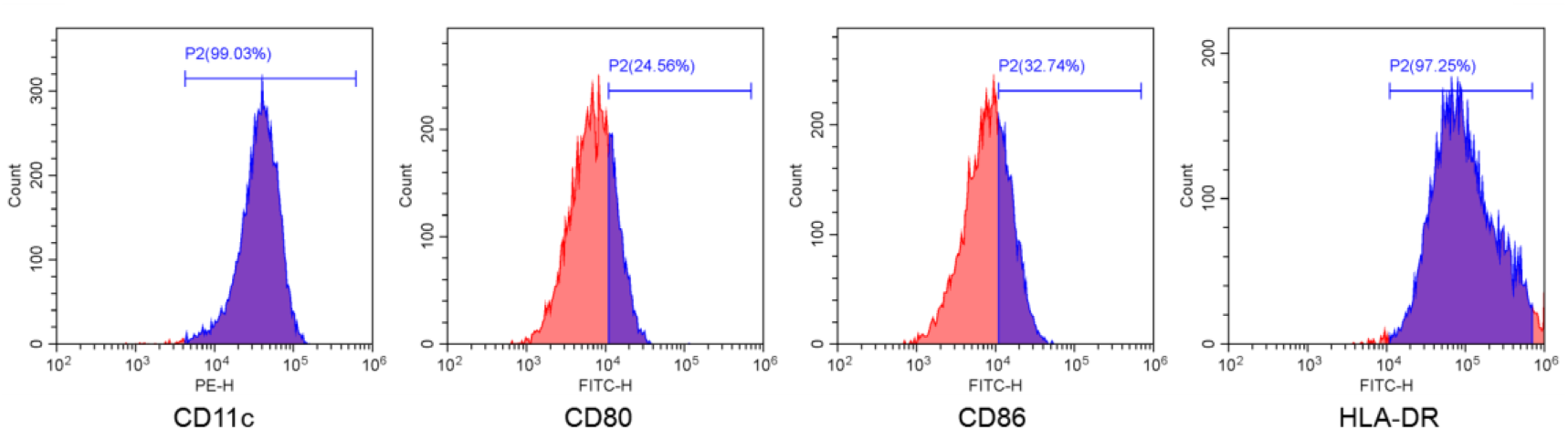
The moDC Markers are Performed by FCM.

### moDC morphological and functional differences

Scanning electron microscopy showed that moDC of UA, NST, and ST groups had longer and denser pseudopod-like processes than that of CON group in differences of morphology(Figure 5).

**Figure 5.**
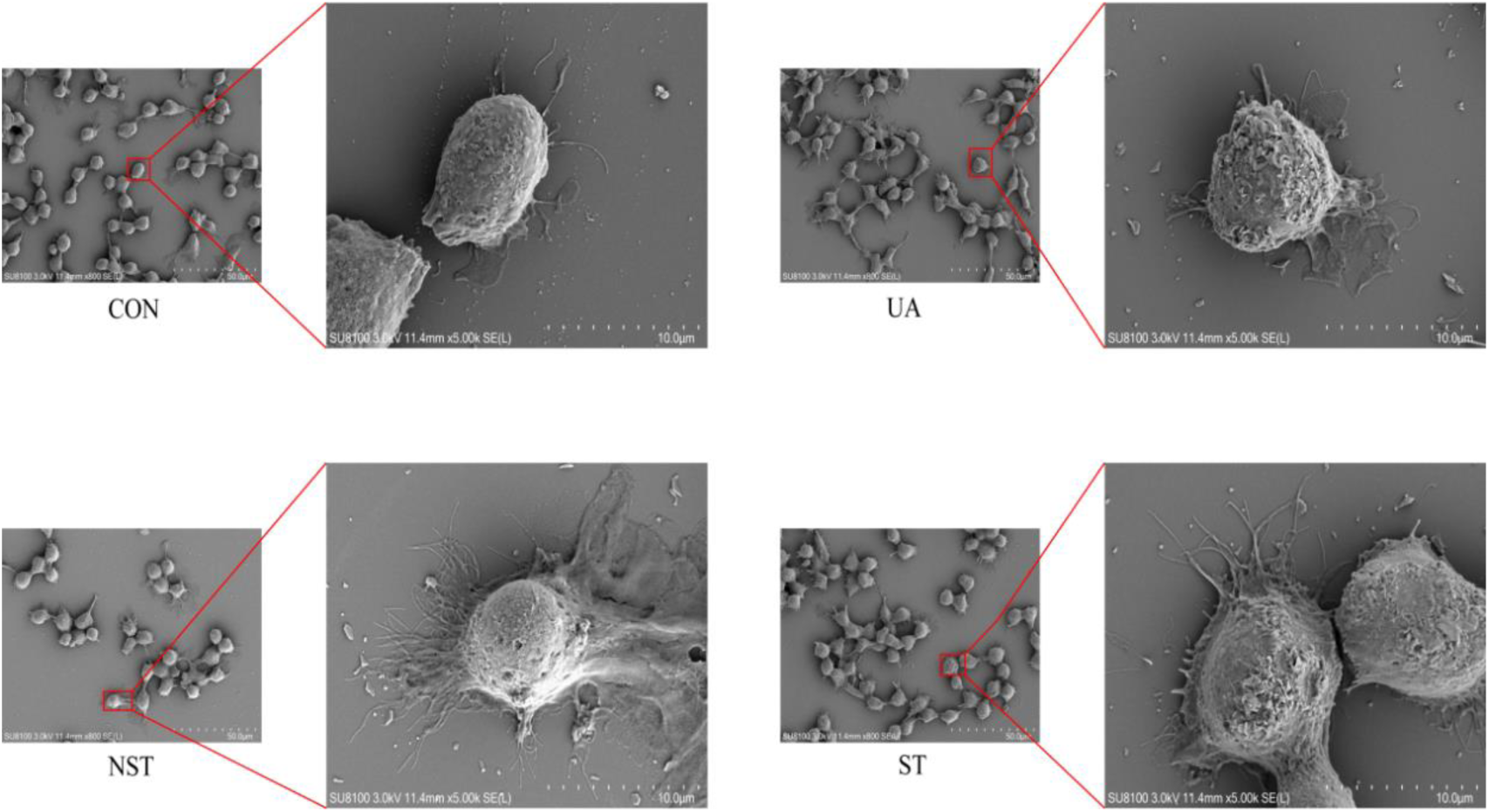
Morphological Differences of moDC among Different Groups under Scanning Electron Microscopy.

The results of FCM showed that the expression levels of CD80 and HLA-DR and mean fluorescence intensity (MFI) in ST and NST groups were higher than those in CON and UA groups(p<0.05), while there was no statistical difference in the expression levels of CD80 and HLA-DR between CON and UA groups, NST and ST groups in antigen presentation function of moDC(Figure 6), and the expression level and MFI value of moDC OVA in ST group were lower than those in CON and UA groups (p<0.05), while MFI value showed no statistical difference between NST group, CON group and UA group in antigen presentation and phagocytosis function of moDC(Figure 7). The results of ELISA showed that the expression levels of IL-6 and IL-12p70 in ST and NST groups were higher than those in CON and UA groups (p<0.05) (Figure 8A, C), while there was no statistical difference in IL-10 expression levels among all groups in secretory function of moDC(Figure 8B).

**Figure 6.**
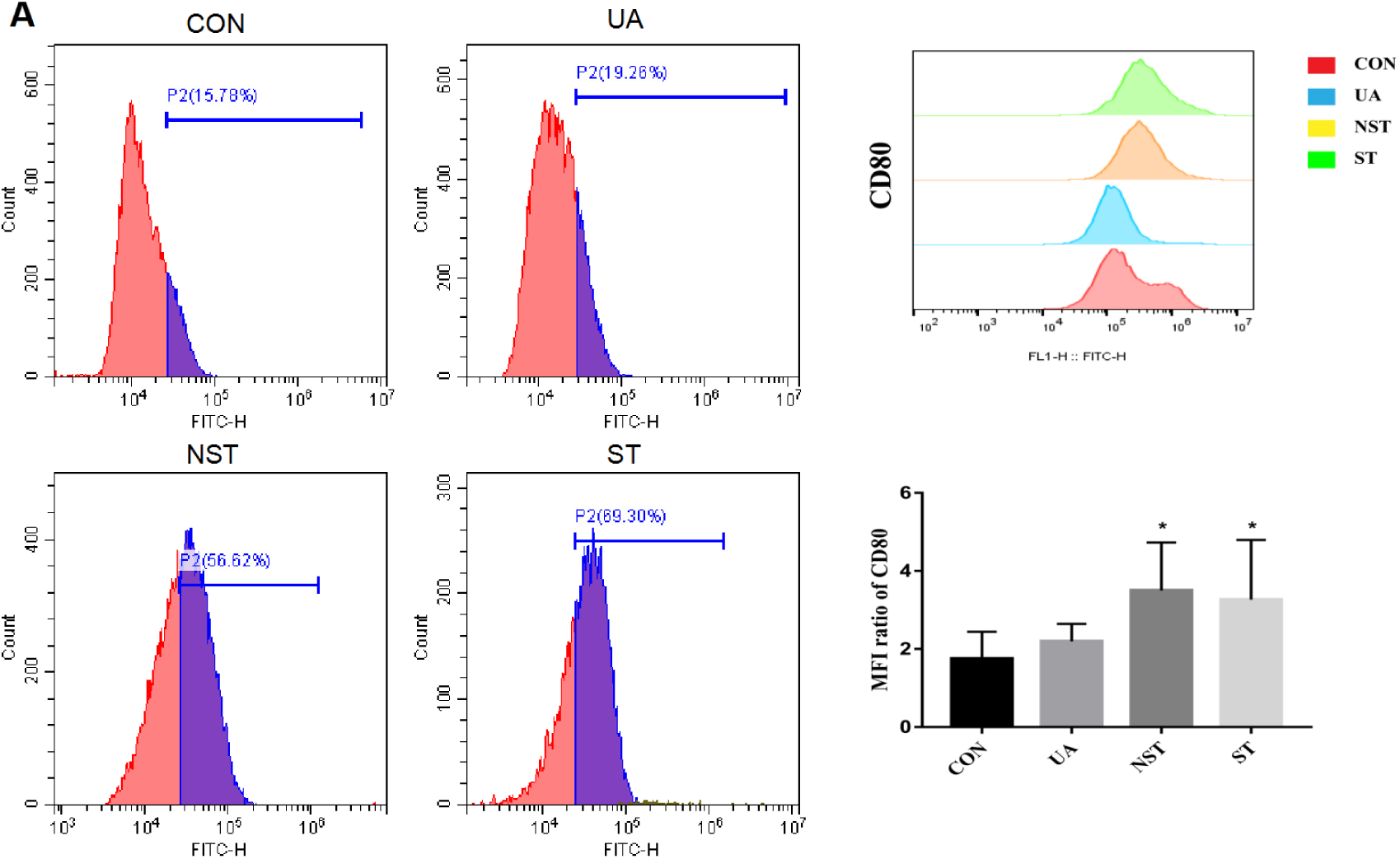
The Differences of moDC Antigen Presentation among Different Groups by FCM.

**Figure 7.**
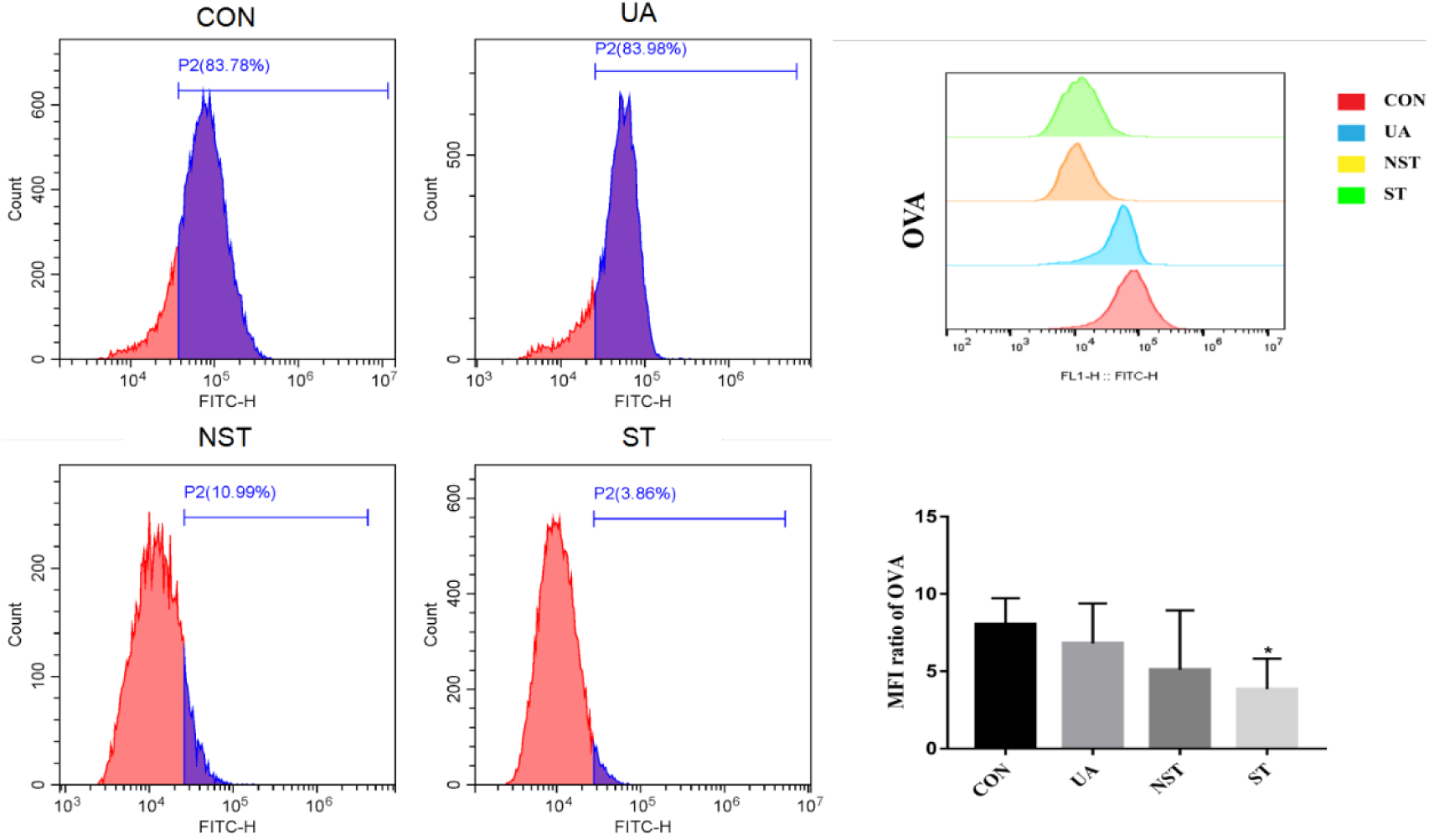
The differences of moDC Phagocytosis among Different Groups by FCM.

**Figure 8.**
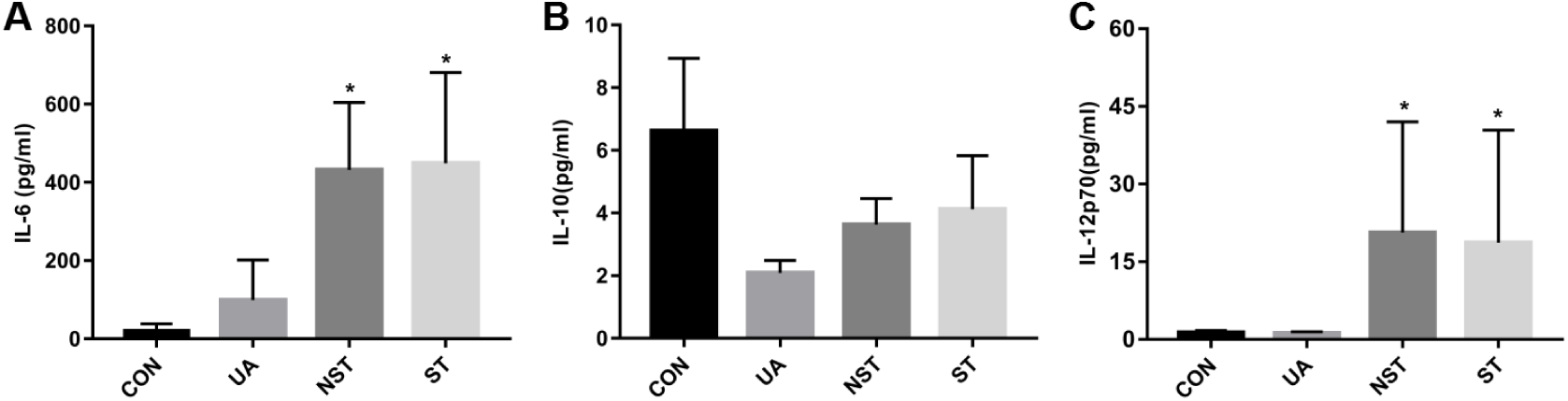
The Differences of moDC Secretion among Different Groups by FCM. Note: *p<0.05, compared with CON.

### Differential expression of lncRNA

The results of sequencing showed that CON vs UA, 3 lncRNA: 1 up-regulated and 2 down-regulated; CON vs NST, 49 lncRNA: 13 up-regulated and 36 down-regulated; CON vs ST, 35 lncRNA:18 up-regulated and 17 down-regulated; UA vs NST, 115 lncRNA:44 up-regulated and 71 down-regulated. UA vs ST, 113 lncRNA: 70 up-regulated and 43 down-regulated; ST vs NST, 4 lncRNA:0 up-regulated and 4 down-regulated (Table 2). The top 10 lncRNA were significantly differentially expressed between UA and STEMI patients(Table 3); The top 10 lncRNA were significantly differentially expressed between healthy subjects and STEMI patients (Table 4). The statistics of differentially expressed lncRNA were calculated using volcano maps (Figure 9), and the distribution of which was illustrated by heat map (Figure 10).

**Figure 9.**
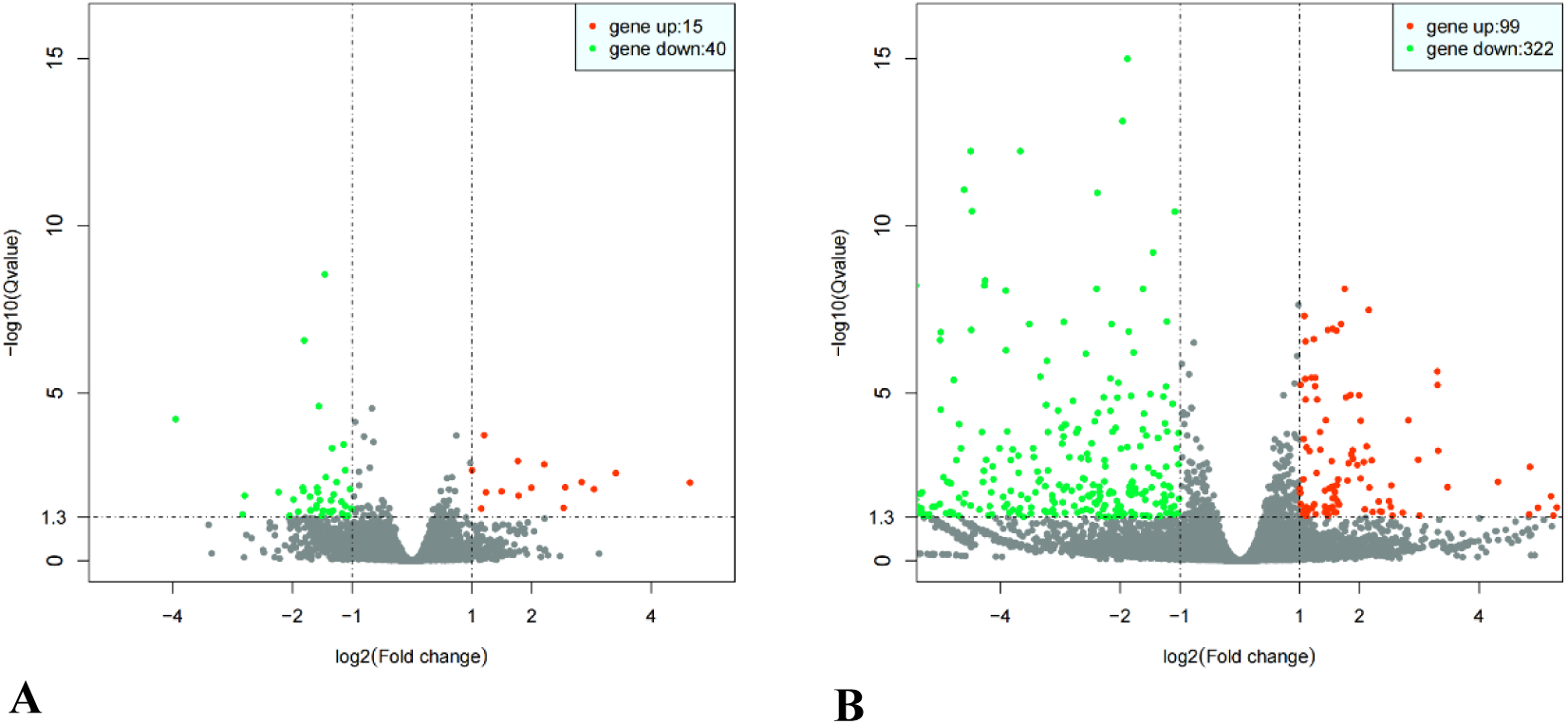
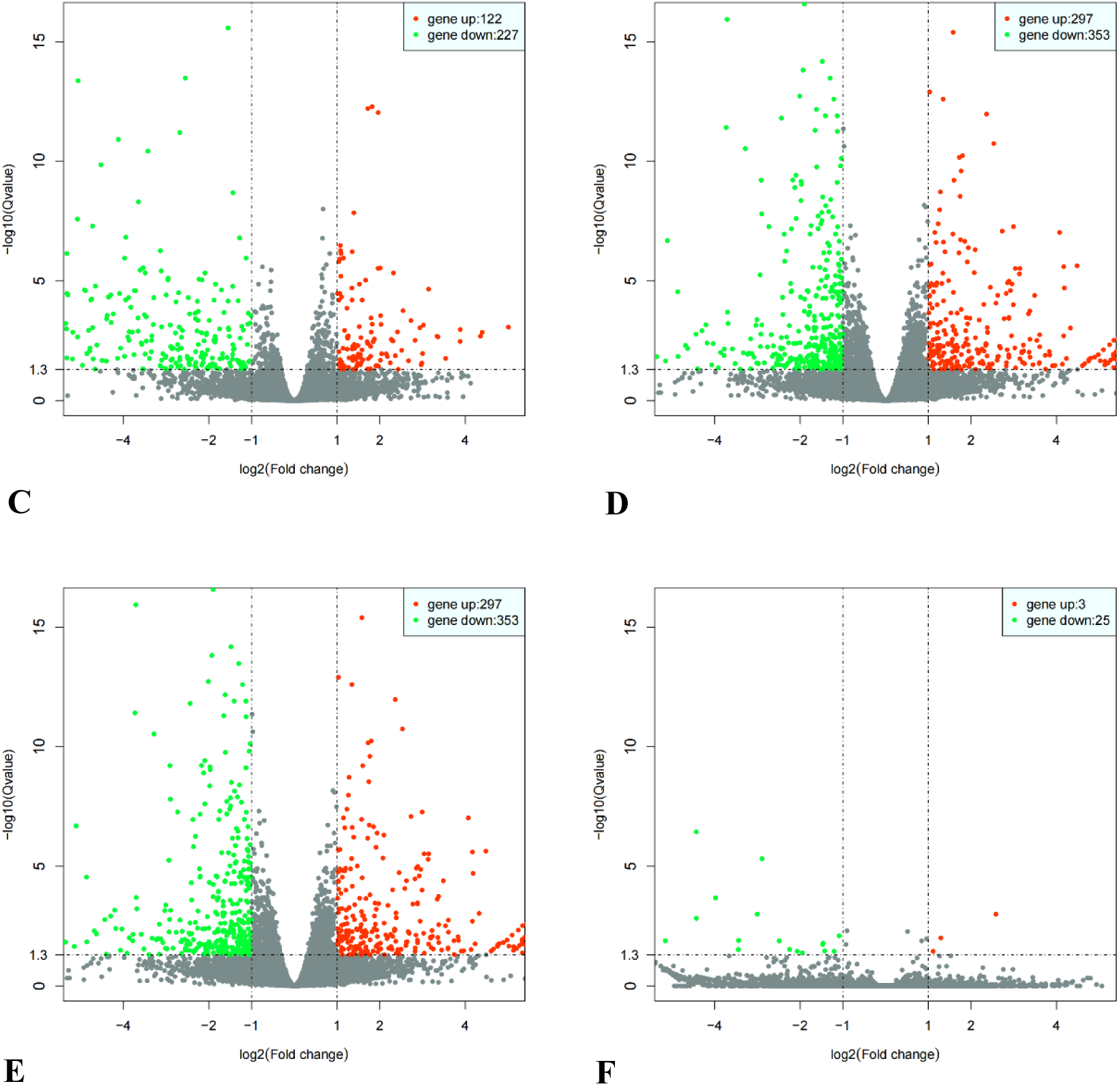
Differential Expression Volcano Plot of lncRNAs and mRNAs in Differen Types of Patients. Note:Red dots represent up regulated lncRNAs and mRNAs, and green dots represent down regulated lncRNAs and mRNAs with statistical significance (fold change≥2, *P* < 0.05). while the gray dots are not statistically significant (*P* > 0.05). A: CON vs UA(n=3); B: CON vs NST(n=3); C:CON vs NST(n=3); D: CON vs NST(n=3); E: CON vs NST(n=3); F: CON vs NST(n=3).

**Figure 10.**
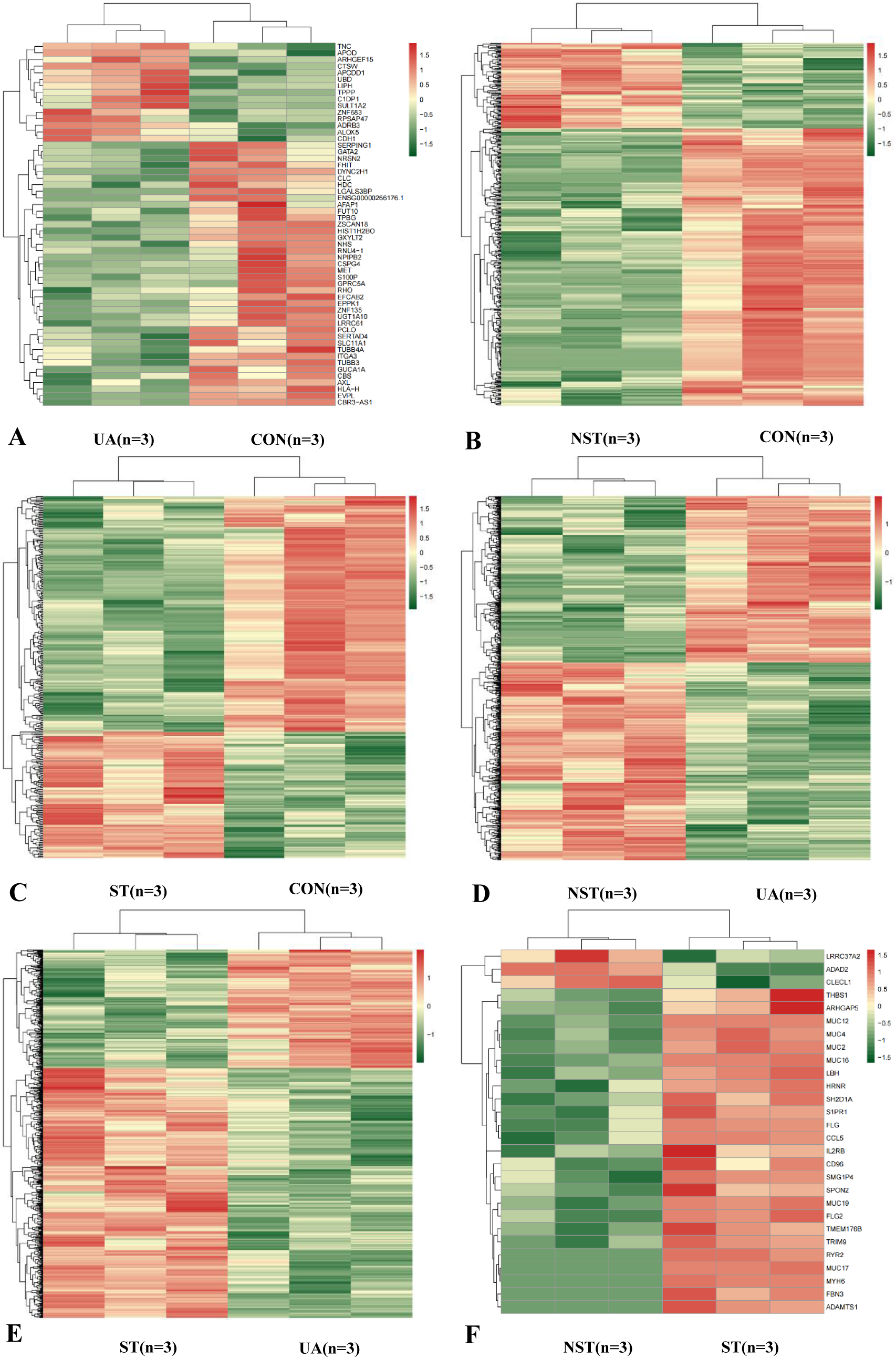
The Differential Expression Heatmap of lncRNA and mRNA in Different Types of Patients. Note:Red color indicates high relative expression and green color indicates low relative expression.

**Table 2.** LncRNAs Expression Differences among Different Types of Patients.

| Compare-group | Sig | Up | Down |
| --- | --- | --- | --- |
| CON-UA | 3 | 1 | 2 |
| CON-NST | 49 | 13 | 36 |
| CON-ST | 35 | 18 | 17 |
| UA-NST | 115 | 44 | 71 |
| UA-ST | 113 | 70 | 43 |
| NST-ST | 4 | 0 | 4 |

Note: Sig= number of significantly differentially expressed genes; UP= significantly up-regulated gene number; Down= Significantly down-regulated number of genes.

**Table 3.** The Top 10 of Differentially Expressed LncRNAs According to the Fold Change (FC) Values in ST Patients Compared with UA Patients.

| Upregulated lncRNAs |  | Down-regulated lncRNAs |  |
| --- | --- | --- | --- |
| lncRNA ID | Fold change | lncRNA ID | Fold change |
| NR_046502.1 | 6.847151558 | NR_001564.2 | -8.666432781 |
| NR_125734.1 | 6.794857735 | NR_003255.2 | -8.599792289 |
| NR_125733.1 | 6.620243097 | ENST00000381108.3 | -6.432797217 |
| NR_125736.1 | 6.38162831 | ENST00000413897.2 | -5.959498631 |
| NR_125735.1 | 6.36601394 | ENST00000447389.1 | -5.866711676 |
| NR_125737.1 | 5.92137718 | NR_001558.3 | -4.907223863 |
| ENST00000521498.1 | 5.705934793 | ENST00000454861.2 | -4.777113613 |
| NR_001543.3 | 5.432879437 | NR_037843.3 | -4.601534508 |
| ENST00000566193.1 | 5.410383608 | NR_121625.1 | -3.936075356 |
| NR_144458.1 | 5.165316333 | ENST00000453666.1 | -3.917072598 |

Note: lncRNAs=long noncoding RNAs, UA=unstable angina, STEMI=ST-elevation myocardial infarction.

**Table 4.** The Top 10 of Differentially Expressed LncRNAs According to the Fold Change (FC) Values in ST Patients Compared with CON Patients.

| Upregulated lncRNAs |  | Down-regulated lncRNAs |  |
| --- | --- | --- | --- |
| lncRNA ID | Fold change | lncRNA ID | Fold change |
| NR_027922.3 | 4.026332123 | NR_001564.2 | -8.11180645 |
| NR_027921.3 | 3.891106436 | NR_003255.2 | -8.045015752 |
| ENST00000514425.1 | 2.314561714 | NR_001558.3 | -6.842515511 |
| ENST00000585945.1 | 1.903064576 | ENST00000433510.2 | -5.982336514 |
| ENST00000586030.1 | 1.851923392 | NR_037843.3 | -5.18496365 |
| ENST00000590780.1 | 1.822037256 | NR_027701.1 | -3.011757335 |
| ENST00000609178.1 | 1.723826313 | ENST00000608596.1 | -2.327906013 |
| ENST00000434683.1 | 1.700832016 | NR_038885.1 | -2.271360898 |
| ENST00000609272.1 | 1.608128612 | ENST00000491934.2 | -2.056528047 |
| ENST00000592117.1 | 1.596146278 | ENST00000602277.1 | -1.716557573 |

Note: lncRNAs, long noncoding RNAs; CON, healthy volunteer; ST, ST-elevation myocardial infarction.

Five lncRNA (ENST00000446952.1, ENST00000457097.1, ENST00000590780.1, NR_026793.1, NR_027922.3,) were selected to perform qRT-PCR tests according to their conservative scores and differential expression multiple. The results showed that the expression levels of NR_026793.1, NR_027922.3, ENST00000446952.1 and ENST00000590780.1 were up-regulated, while the expression level of ENST00000457097.1 was not statistically significant between the two groups (Figure 11).

**Figure 11.**
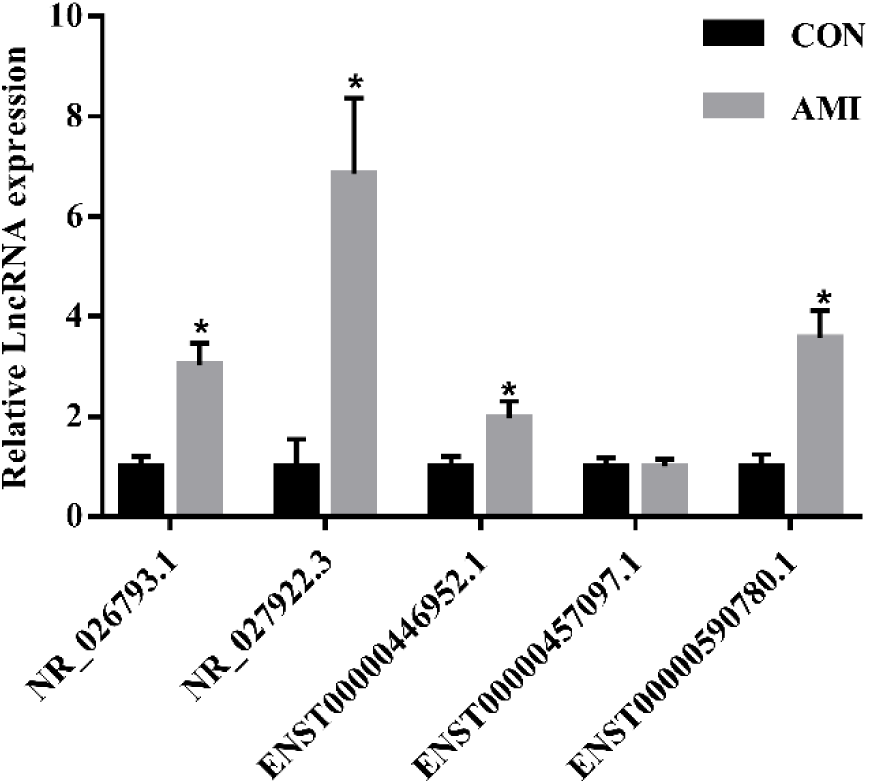
The Sequencing Results were verified by RT-PCR. Note: AMI: n=14,CON: n=5; *p<0.05, compared with CON.

The results of GO analysis showed that the cell components of differentially expressed RNA mainly involved cell, organelle, cell part, etc., the molecular functions mainly involved protein binding, binding, enzyme binding, etc., and biological processes mainly involve single−organism cellular process, cellular process, single−organism process, etc. (Figure 12). According to KEGG’s analysis, the main signaling pathways involved are related to metabolic pathways, steroid hormone biosynthesis, etc. (Figure 13).

**Figure 12.**
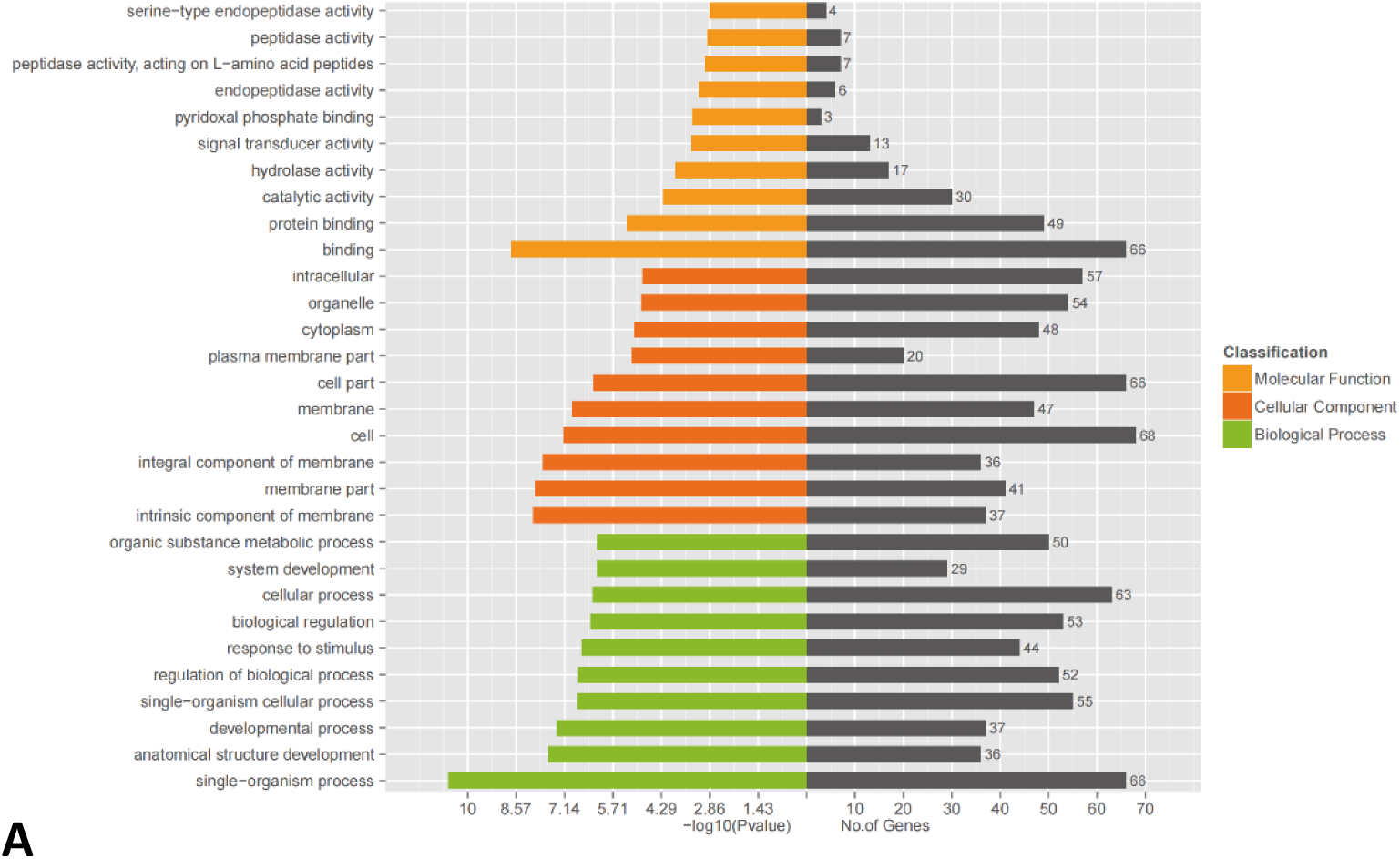
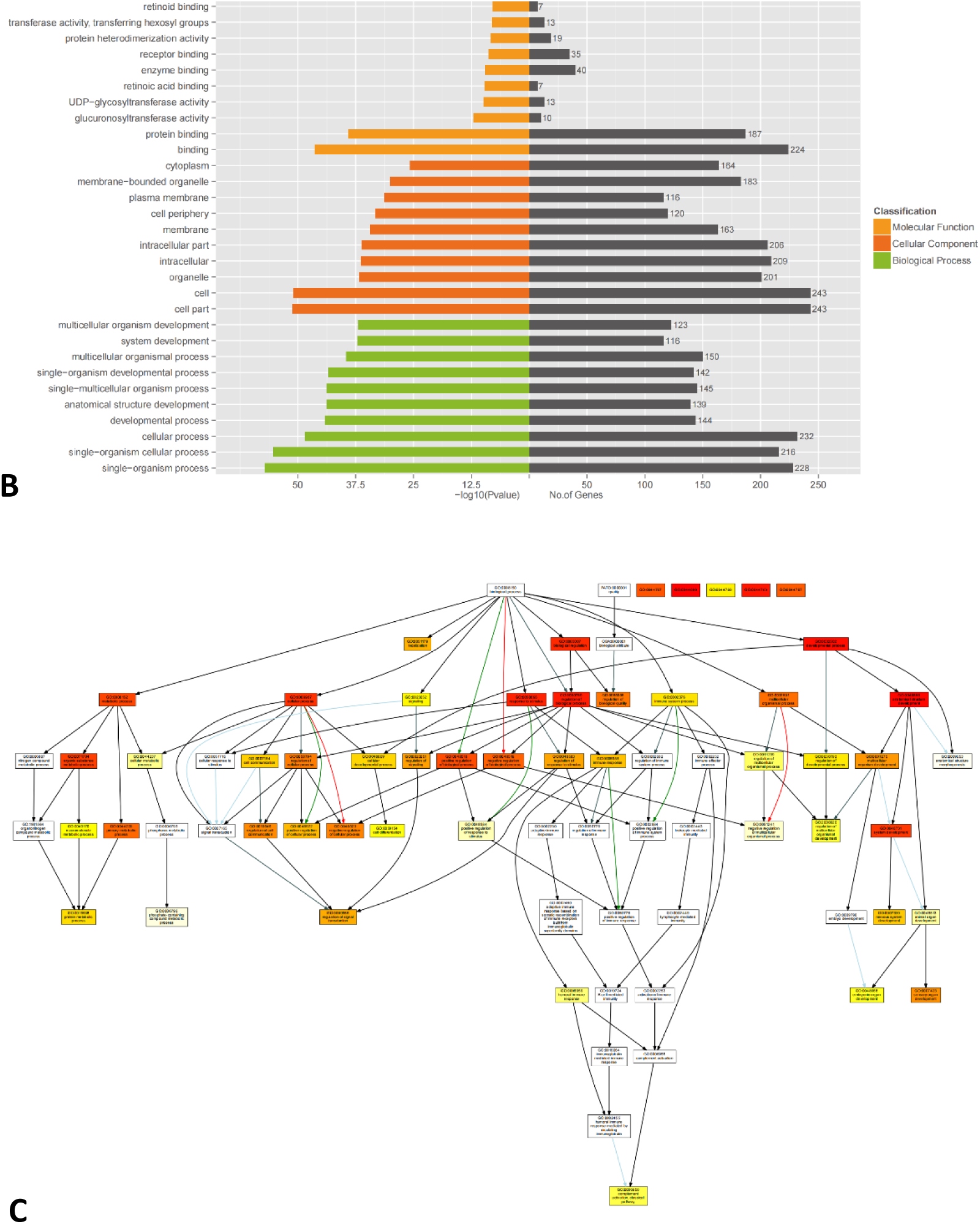
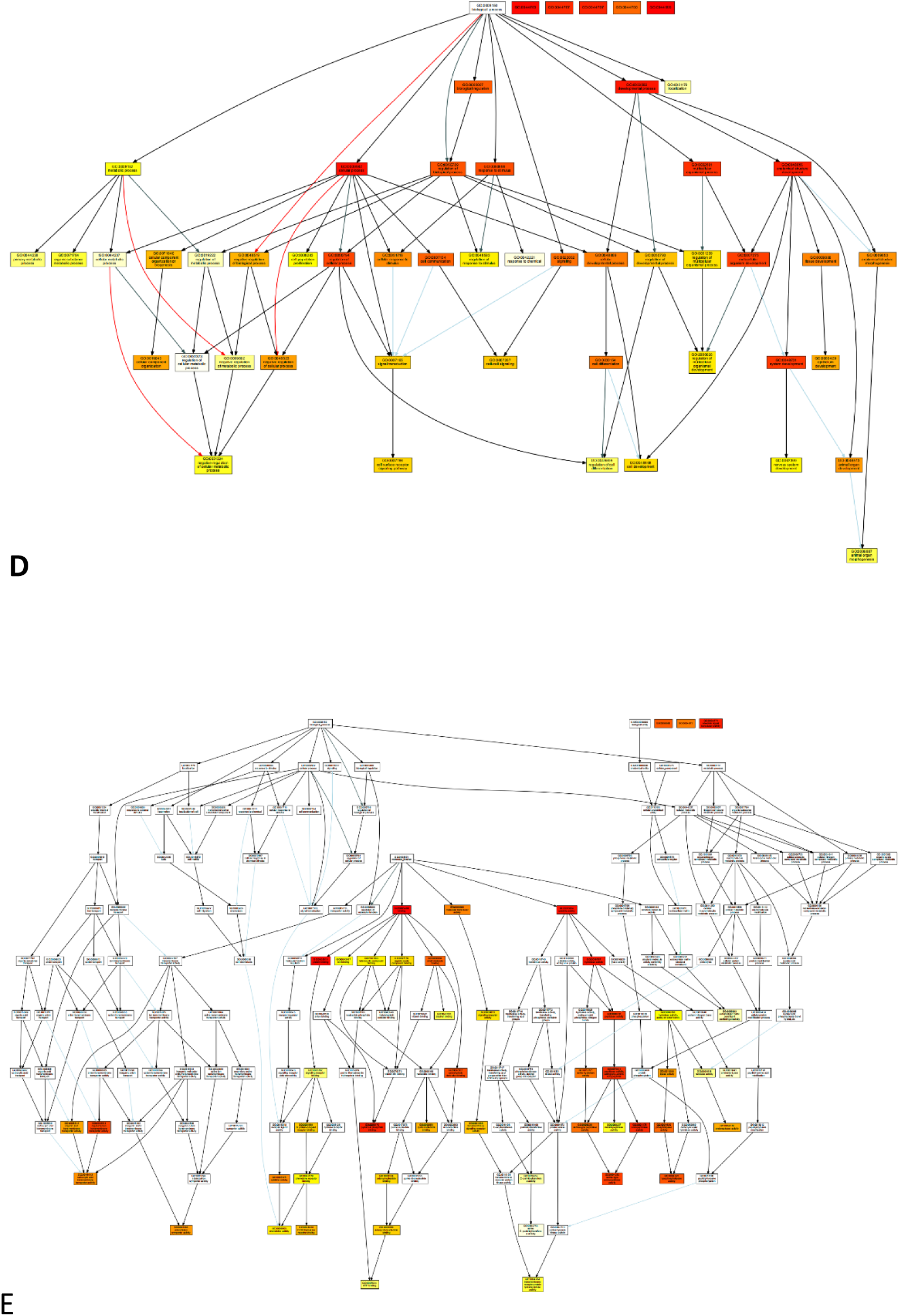
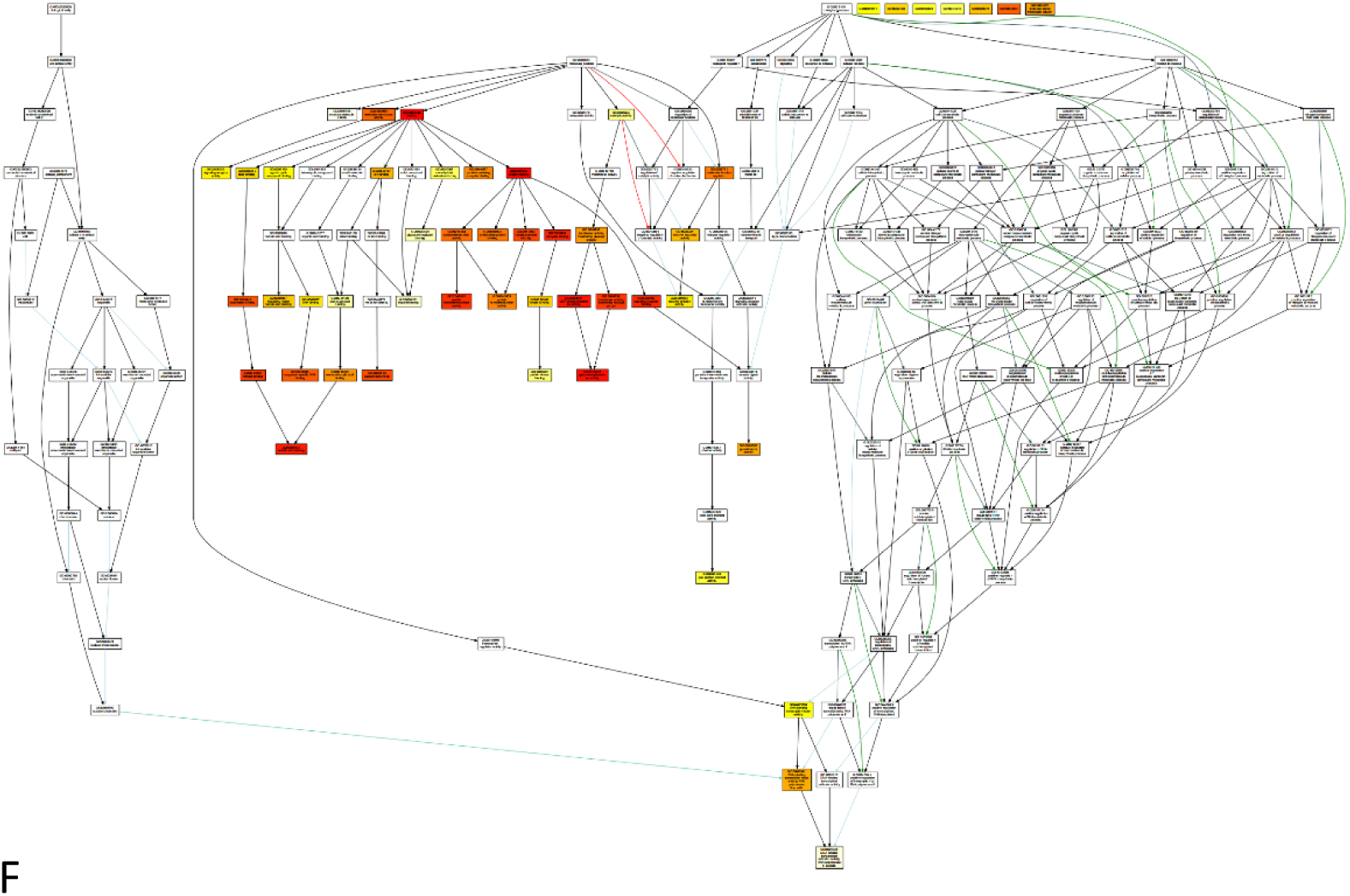
Results of GO Analysis. Notes: A, GO analysis of up regulated RNA; B, GO analysis of down regulated RNA; C, Biological process of up regulated RNA; D, Biological process of down regulated RNA; E, Biological function of up regulated RNA; F, Biological function of down regulated RNA.

**Figure 13.**
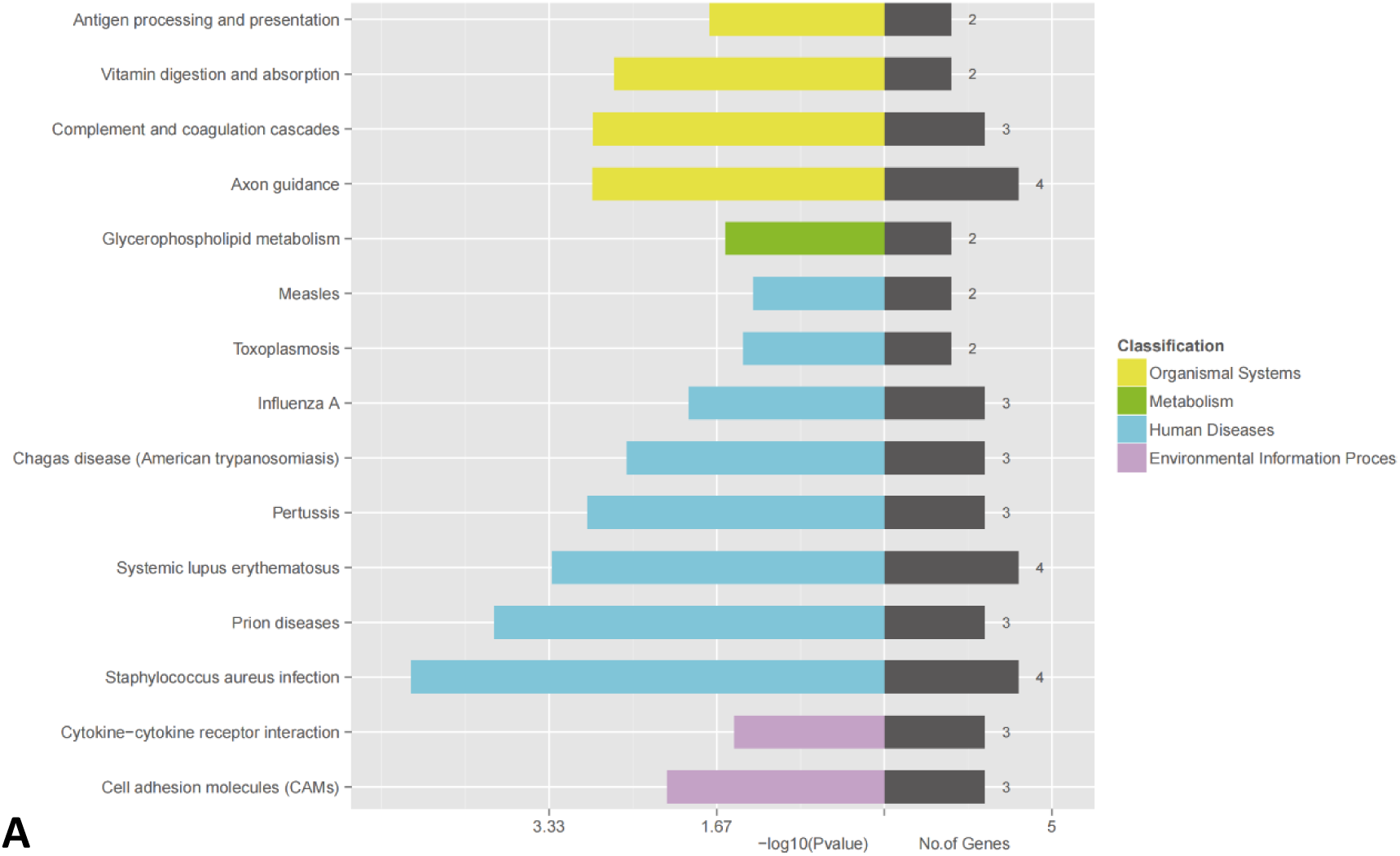
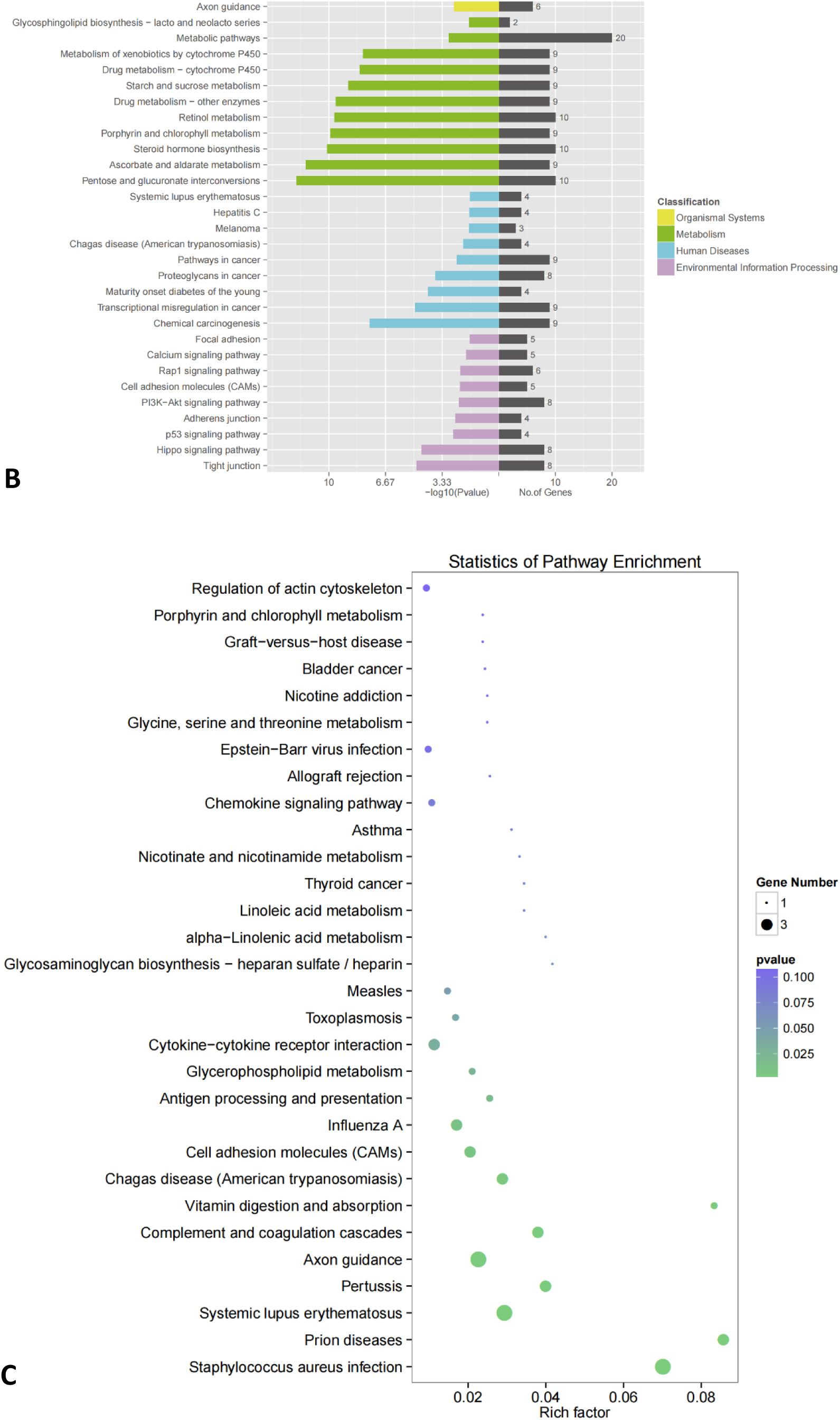
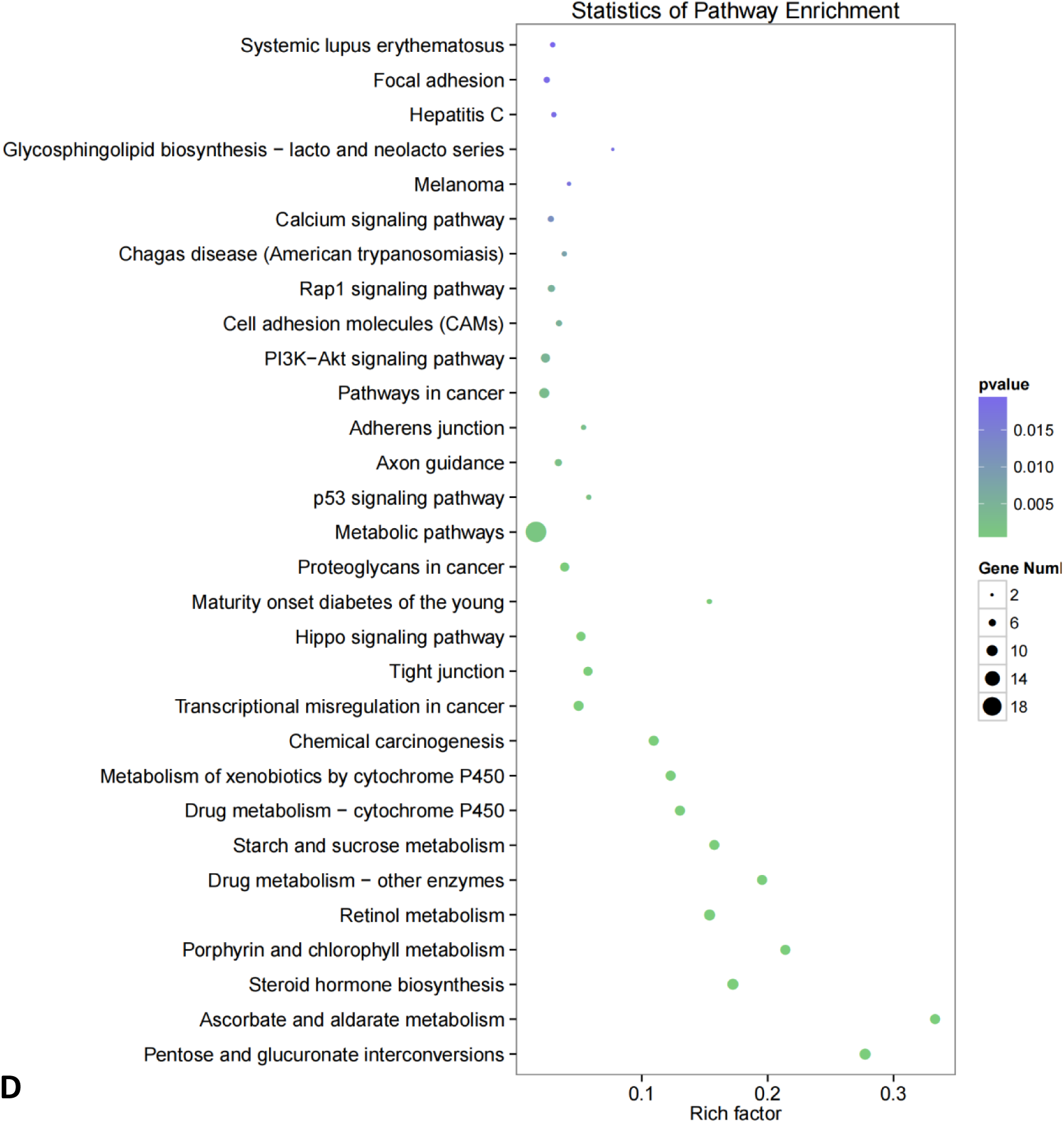
Results of KEGG Analysis. Notes: A, KEGG of up regulated RNA; B, KEGG of down regulated RNA; C, KEGG point of up regulated RNA; D, KEGG point of down regulated RNA.

The results of pearson correlation coefficient showed that ENST00000590780.1 (R=0.61, p<0.05) was positively correlated with CD80 (Figure 14A), ENST00000446952.1 (R=0.61, p<0.05), NR_027922.3 (R=0.56, p<0.05) were positively correlated with CD86 (Figure 14B, C), NR_027922.3 (R=0.47, p<0.05), ENST00000446952.1 (R=0.69, p<0.05) was positively correlated with WBC (Figure 14 D-E), NR_027922.3 (R=0.60, p<0.05), ENST00000446952.1 (R=0.63, p<0.05) was positively correlated with CK-MB (Figure 14F, G), NR_027922.3 (R=0.66, p<0.05), ENST00000446952.1 (R=0.58, p<0.05) was positively correlated with TNT-hs (Figure 14 H, I).

**Figure 14.**
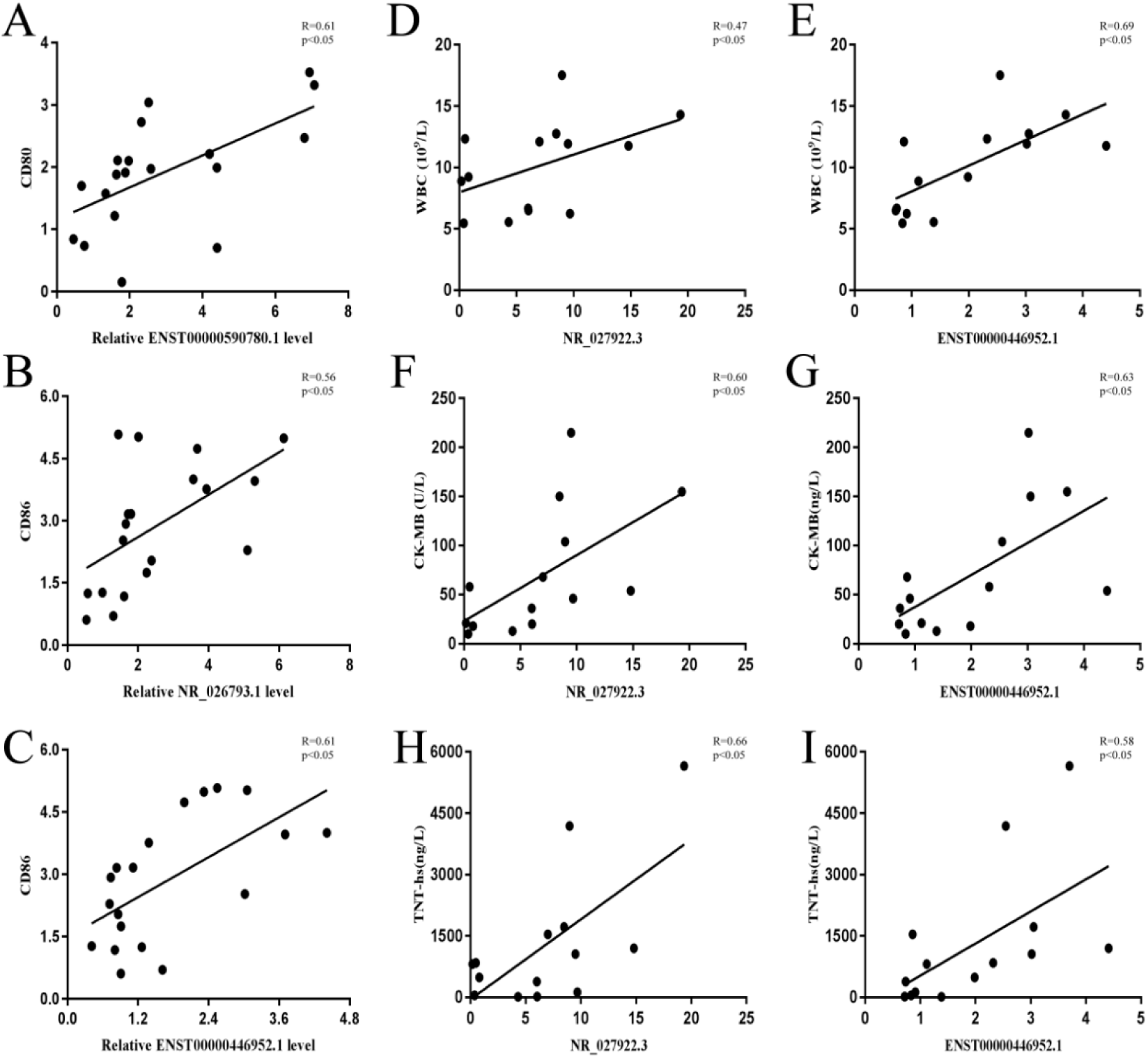
The Correlation between Candidate lncRNA and Target Genes was Performed by Pearson Correlation Coefficient. Note: AMI: n=14,CON: n=5; *p<0.05, compared with CON.

### The role and mechanism of lncRNA CCL15-CCL14

#### (1) Lentiviral transfection

The results of immunofluorescence showed that MOI=60 and infection time =48h were the best conditions for lentiviral moDC transfection (Figure 15). The results of amplification and solution curves of CCL15-CCL14 in (Figure 16A-C) and RT-PCR showed that, compared with CON group (imDC) and lentiviral empty vector group (imDC-vector), over-expression of lentiviral vector could up-regulate the expression level of CCL15-CCL14 (p<0.05) (Figure 16D).

**Figure 15.**
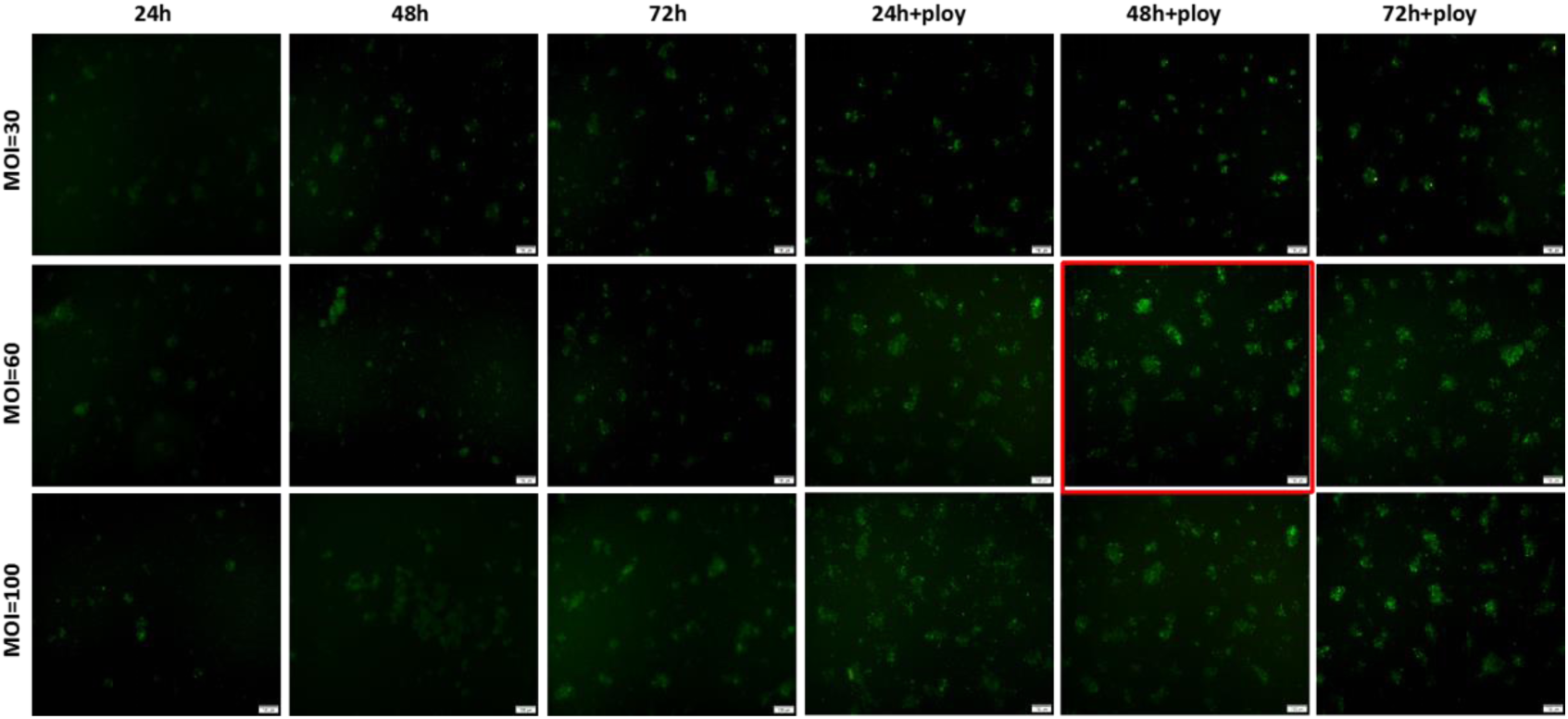
The Fluorescence Intensity of moDC was Observed by Inverted Fluorescence Microscope. Note: Scale =100μm.

**Figure 16.**
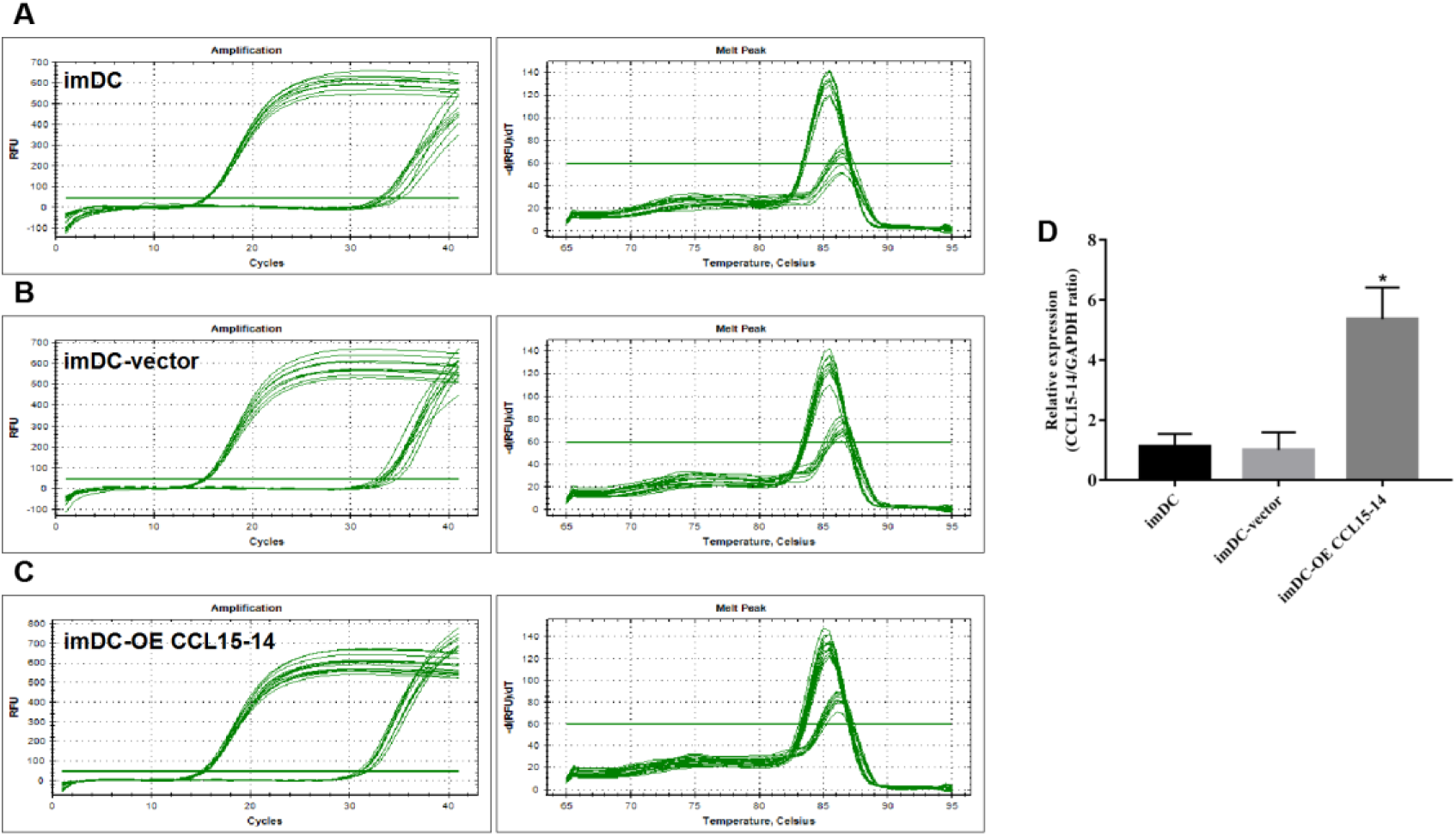
The Amplification and Dissolution Curves and Expression Levels by RT-PCR verified of CCL15-CCL14 after lentivirus over-expression. Note: imDC, immature dendritic cells; imDC-vector, immature dendritic cells-vector; imDC-OE, immature dendritic cells over-expression of lentivirus; n=3 *p<0.05, compared with imDC group.

#### (1) The effect of CCL15-CCL14 transfection

RT-PCR showed that the expression level of CCL15-CCL14 in mature dendritic cells(mDC) increased (p<0.05) (Figure 17); The results of fluorescence microscopy and RT-PCR showed that the expression level of CCL15-CCL14 in moDC decreased after transfection with CCL15-CCL14(Figure 18).

**Figure 17.**
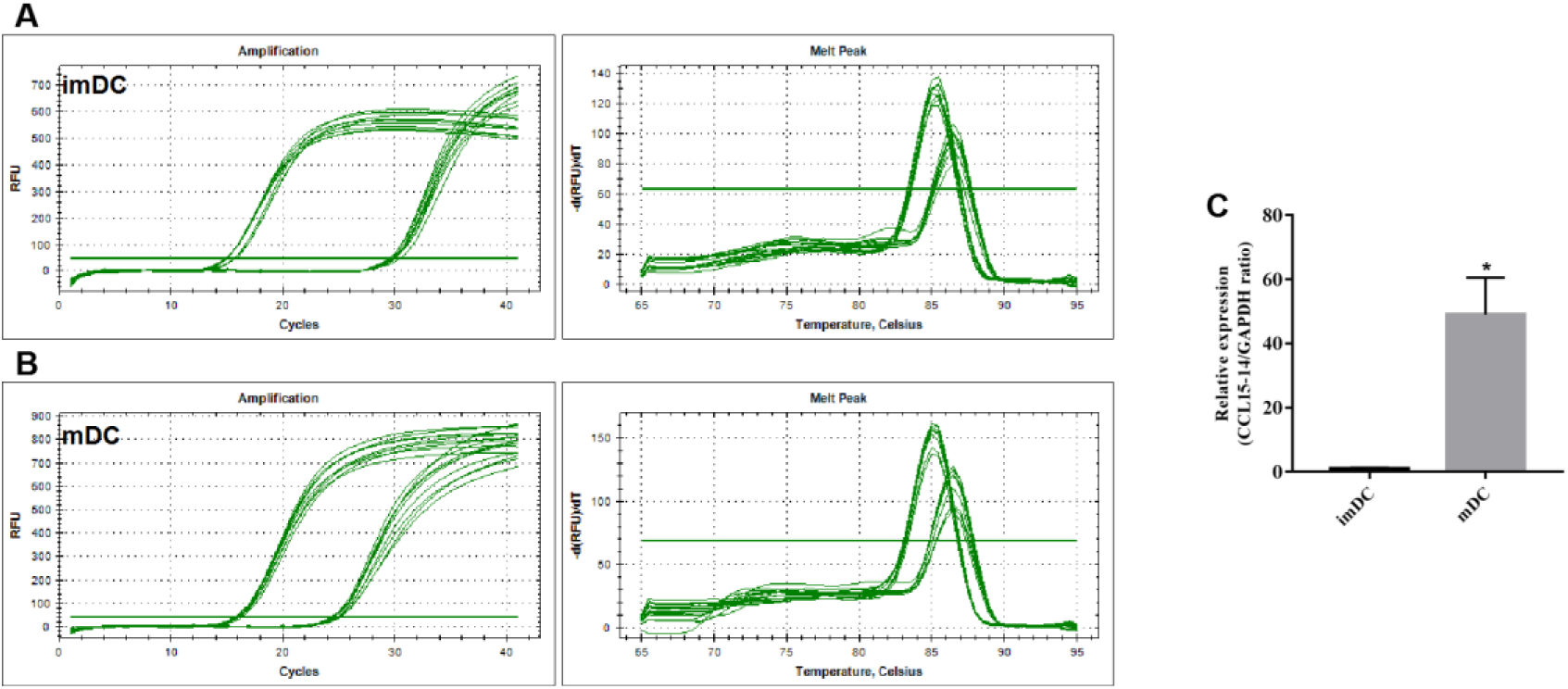
The Amplification and Dissolution Curves and Expression Levels by RT-PCR verified of CCL15-CCL14 after Smart Silencer-CCL15-CCL14. Note: imDC, immature dendritic cells; mDC-vector, mature dendritic cells; n=3 *p<0.05, compared with imDC group.

**Figure 18.**
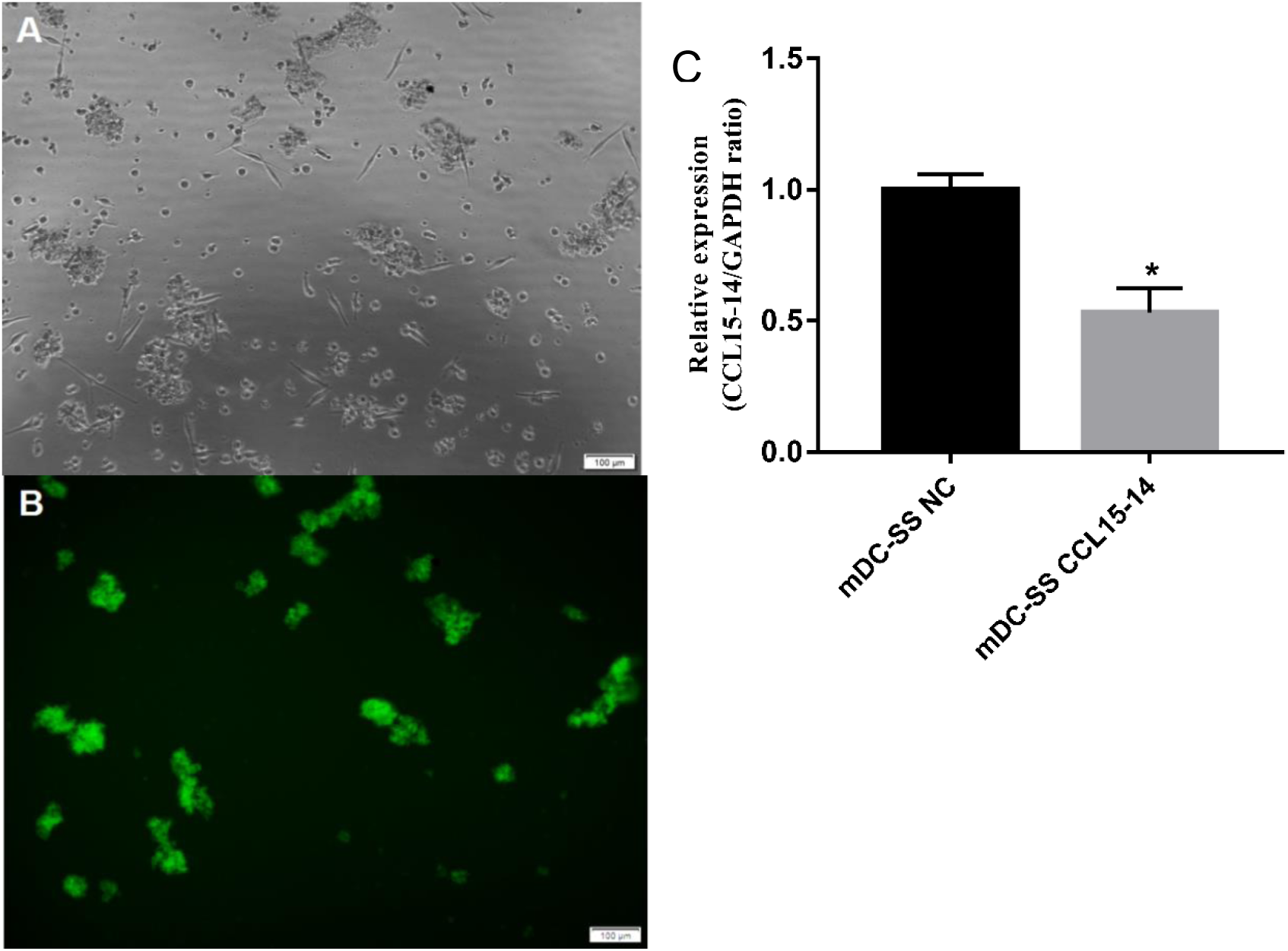
The Transfection Effect of siRNA was Observed by Inverted Fluorescence Microscope and RT-PCR. Note: A and B, results inverted fluorescence microscope; C, results of RT-PCR; Scale =100μm.

#### (2) The effect of CCL15-CCL14 on DC

RT-PCR and FCM showed that the expression levels of CD80, CD86 and HLA-DR increased after over-expression of CCL15-CCL14, conversely, the expression levels of CD80, CD86 and HLA-DR decreased after CCL15-CCL14 silencing (Figure 19 and Figure 20); And the results of ELISA showed that the expression levels of IL-6 and IL-12p70 were increased after over-expression of CCL15-CCL14, while the expression levels of IL-10 were decreased. In contrast, the expression levels of IL-6 and IL-12p70 decreased, and the expression levels of IL-10 increased after CCL15-CCL14 silencing (Figure 21).

**Figure 19.**
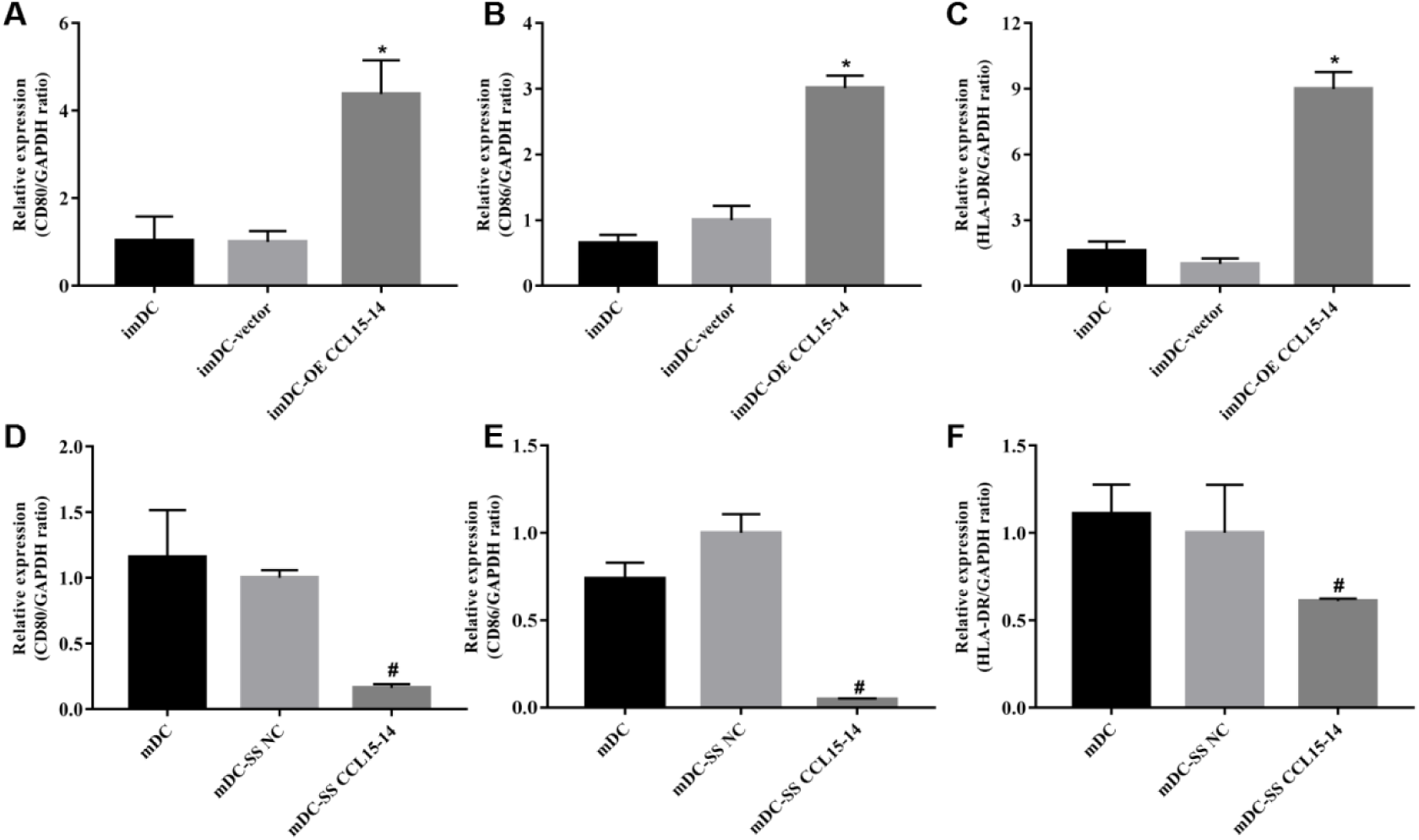
The Expression Level of Markers on DC Surface was Performed by RT-PCR. Note: n=3 *p<0.05, compared with imDC group, #p<0.05, compared with mDC group.

**Figure 20.**
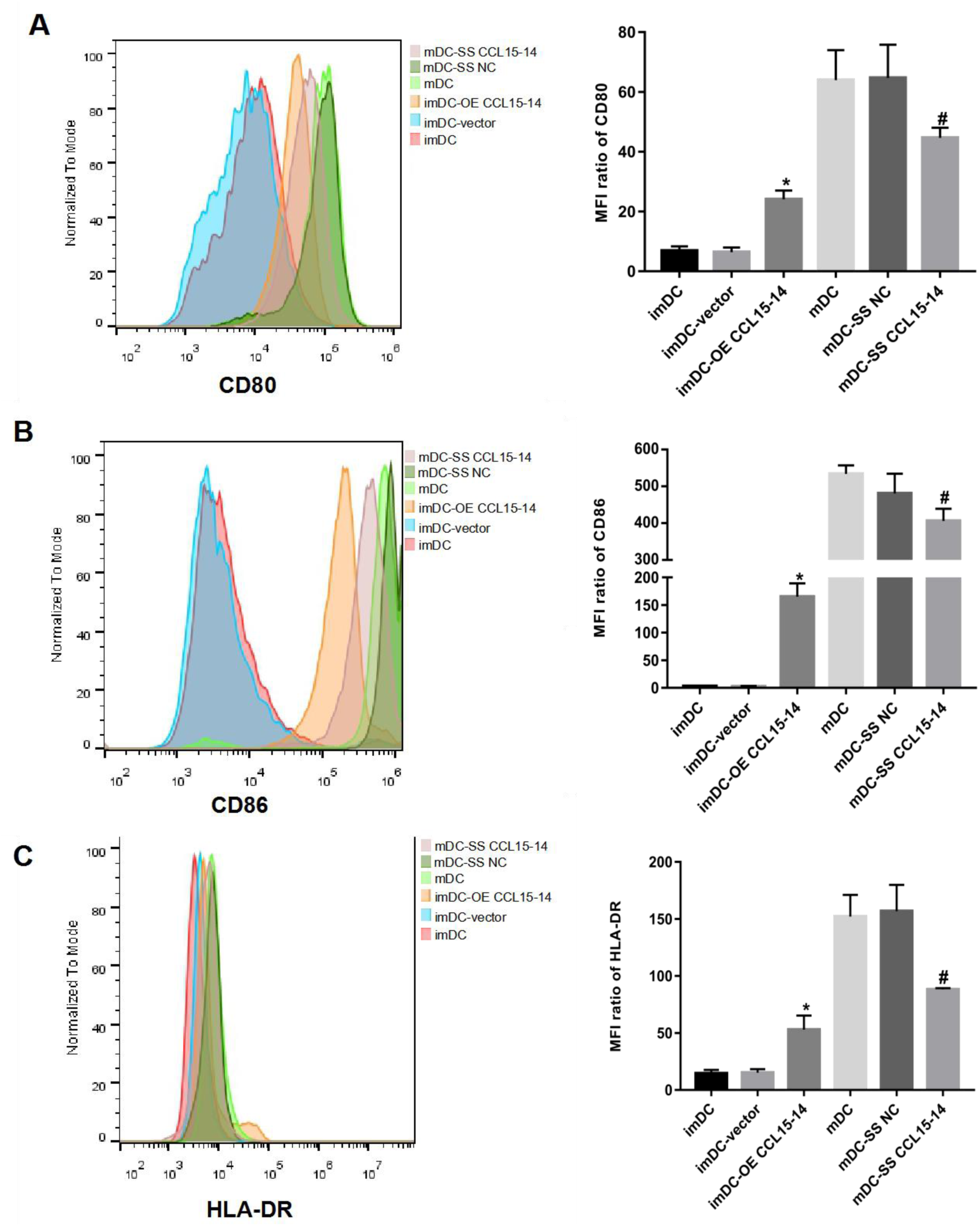
The Expression Level of Markers on DC Surface was Performed by FCM. Note: n=3 *p<0.05, compared with imDC group, #p<0.05, compared with mDC group.

**Figure 21.**
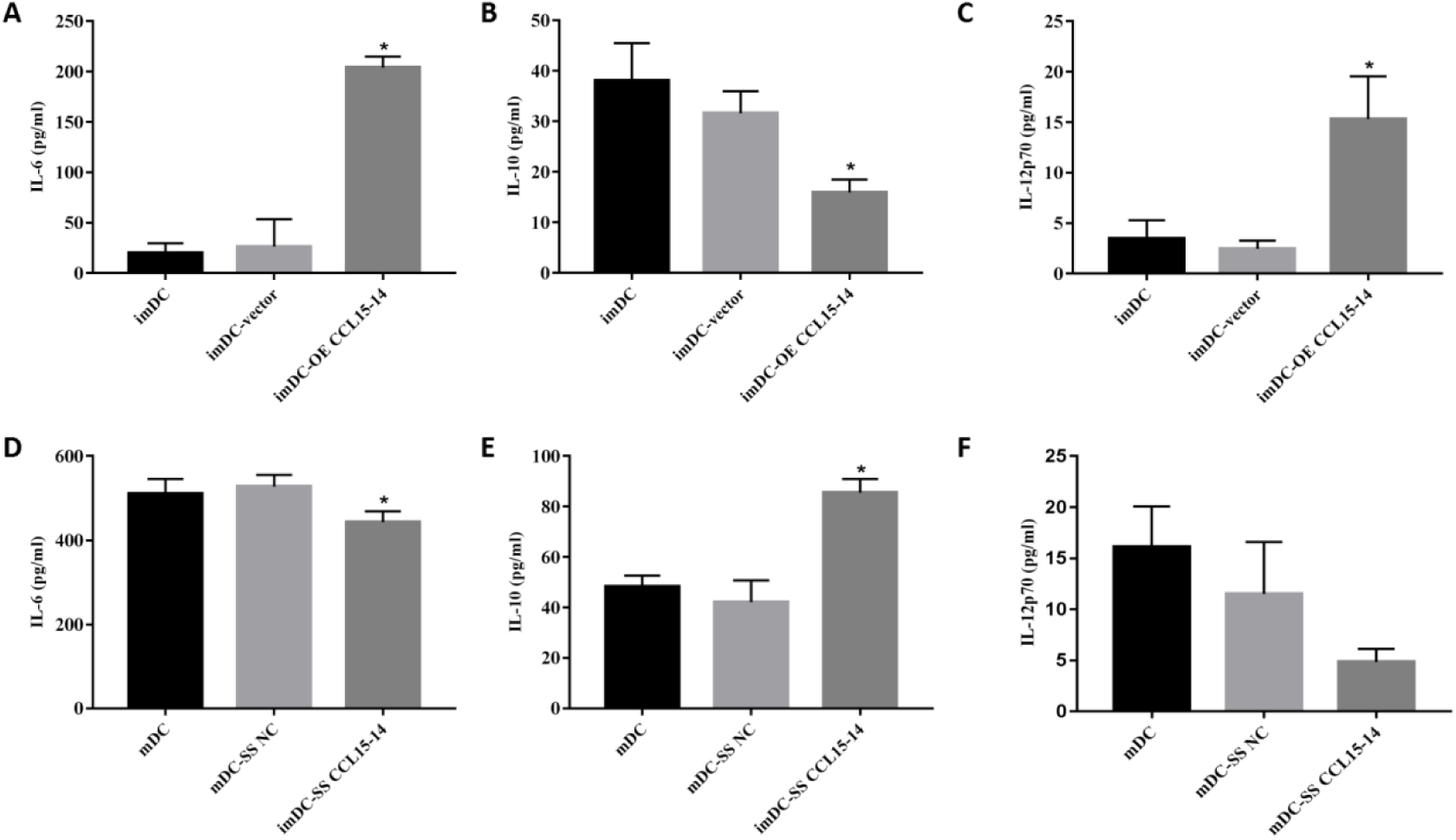
The Secretory Function of DC was Performed by ELISA Methods. Note: n=3 *p<0.05, compared with imDC/mDC group.

#### (3) Pathway proteins of lncRNA

The results of WB showed that overexpression of CCL15-CCL14 could up-regulate the expression levels of p-PI3K and p-AKT, while silencing CCL15-CCL14 could down-regulate the expression levels of p-PI3K and p-AKT (Figure 22).

**Figure 22.**
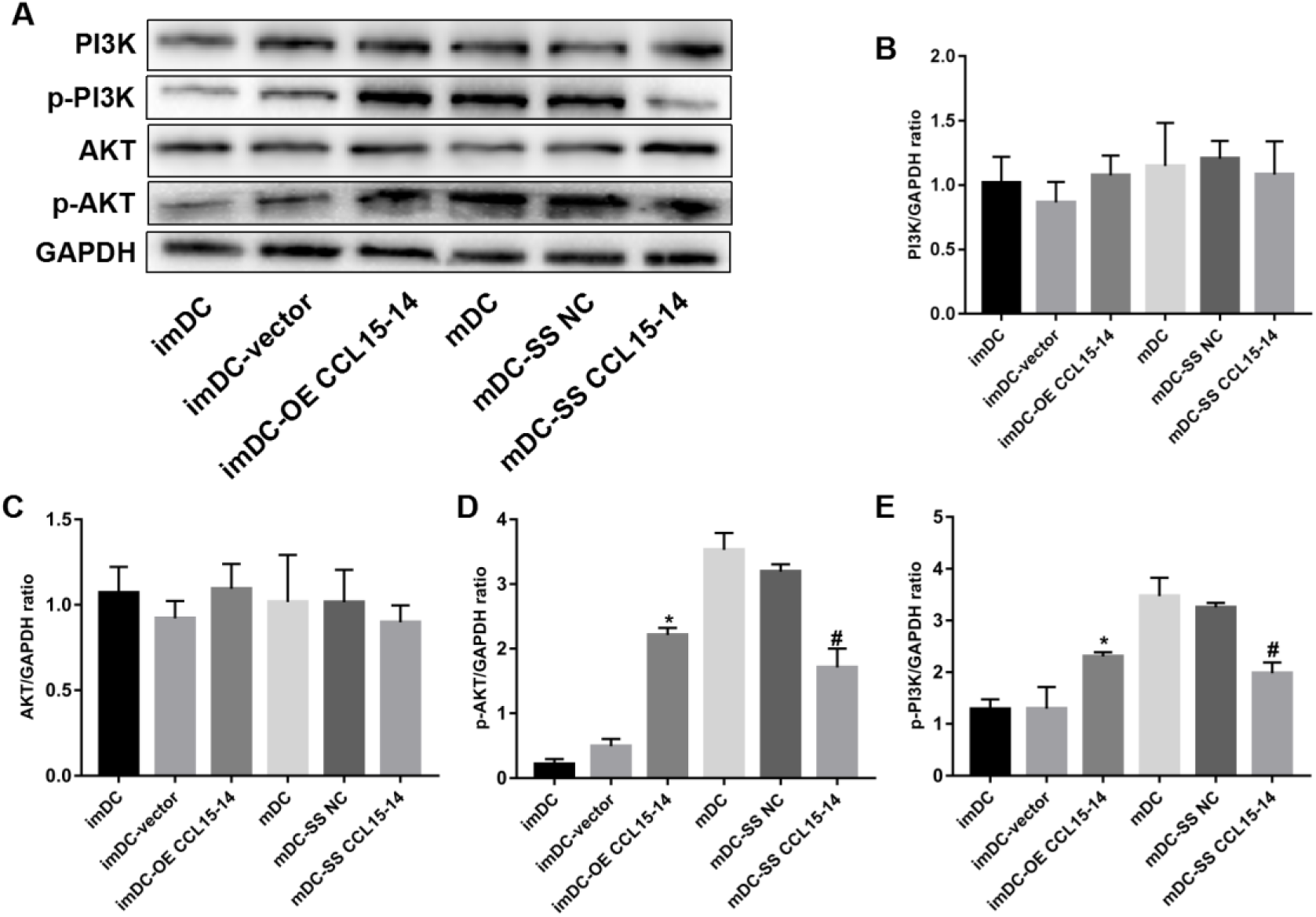
The Potential Pathways of CCL15-CCL14 Function was Performed by Western Blot. Note: n=3 *p<0.05, compared with imDC group, #p<0.05, compared with mDC group.

#### (4) lncRNA subcellular localization

The results of fluorescence in situ hybridization showed that CCL15-CCL14 was mostly expressed in the nucleus of moDC (Figure 23).

**Figure 23.**
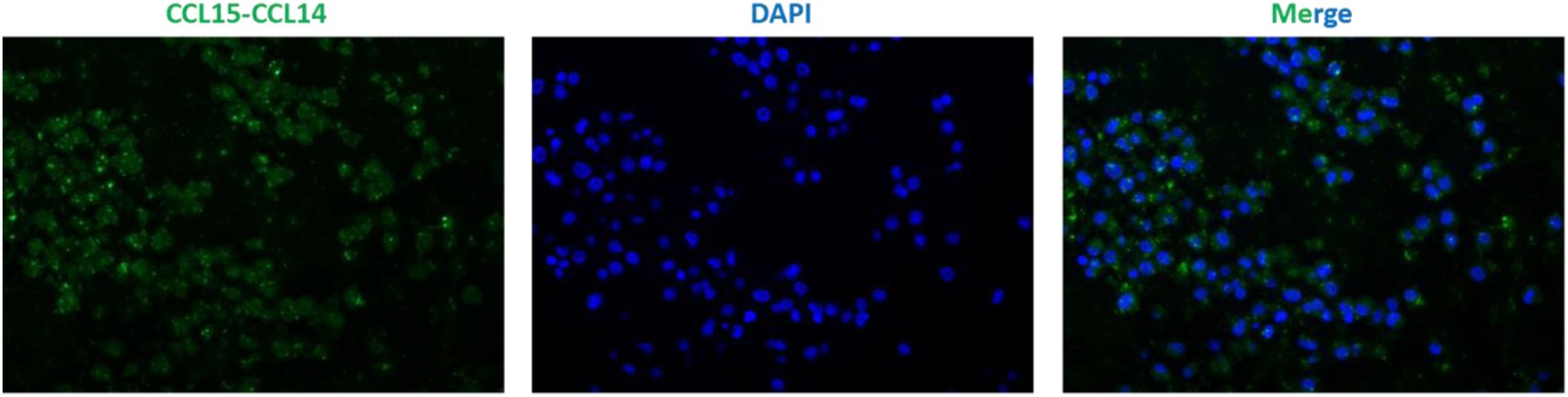
The Subcellular Localization of CCL15-CCL14 was Performed by Immunofluorescence in situ Hybridization. Note: Scale =100μm.

#### (5) The regulatory effect of lncRNA CCL15-CCL14 on CCL14

The results of RT-PCR, ELISA and WB showed that over-expression of CCL15-CCL14 could up-regulate the expression level of CCL14. On the contrary, silencing CCL15-CCL14 could down-regulate the expression level of CCL14. However, the expression level of CCL15 did not change with the expression level of lncRNA CCL15-CCL14(Figure 24-Figure 26).

**Figure 24.**
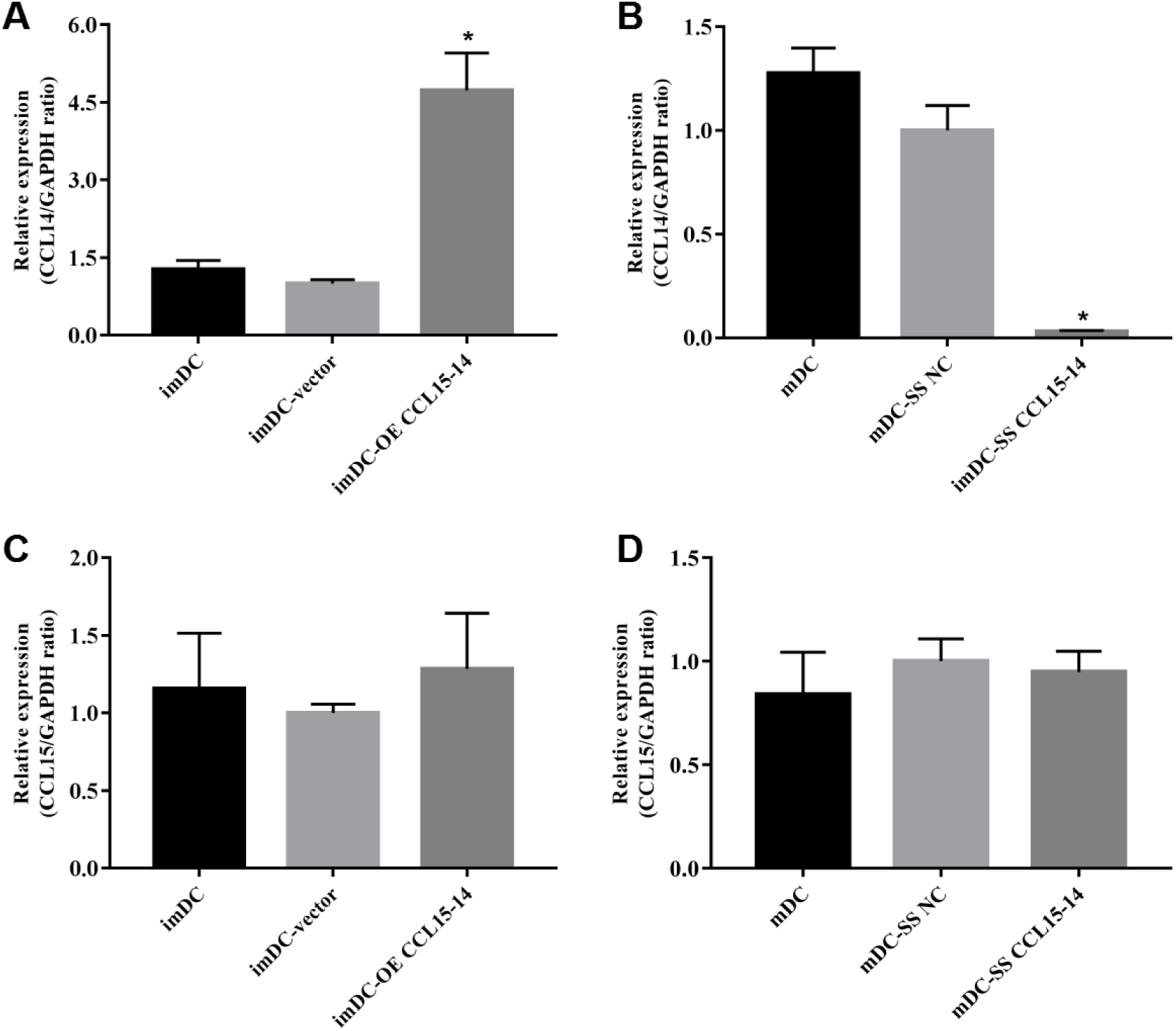
The Expression Levels of moDC CCL14 and CCL15 in each group were Performed by RT-PCR. Note: n=3 *p<0.05, compared with imDC/mDC group.

**Figure 25.**
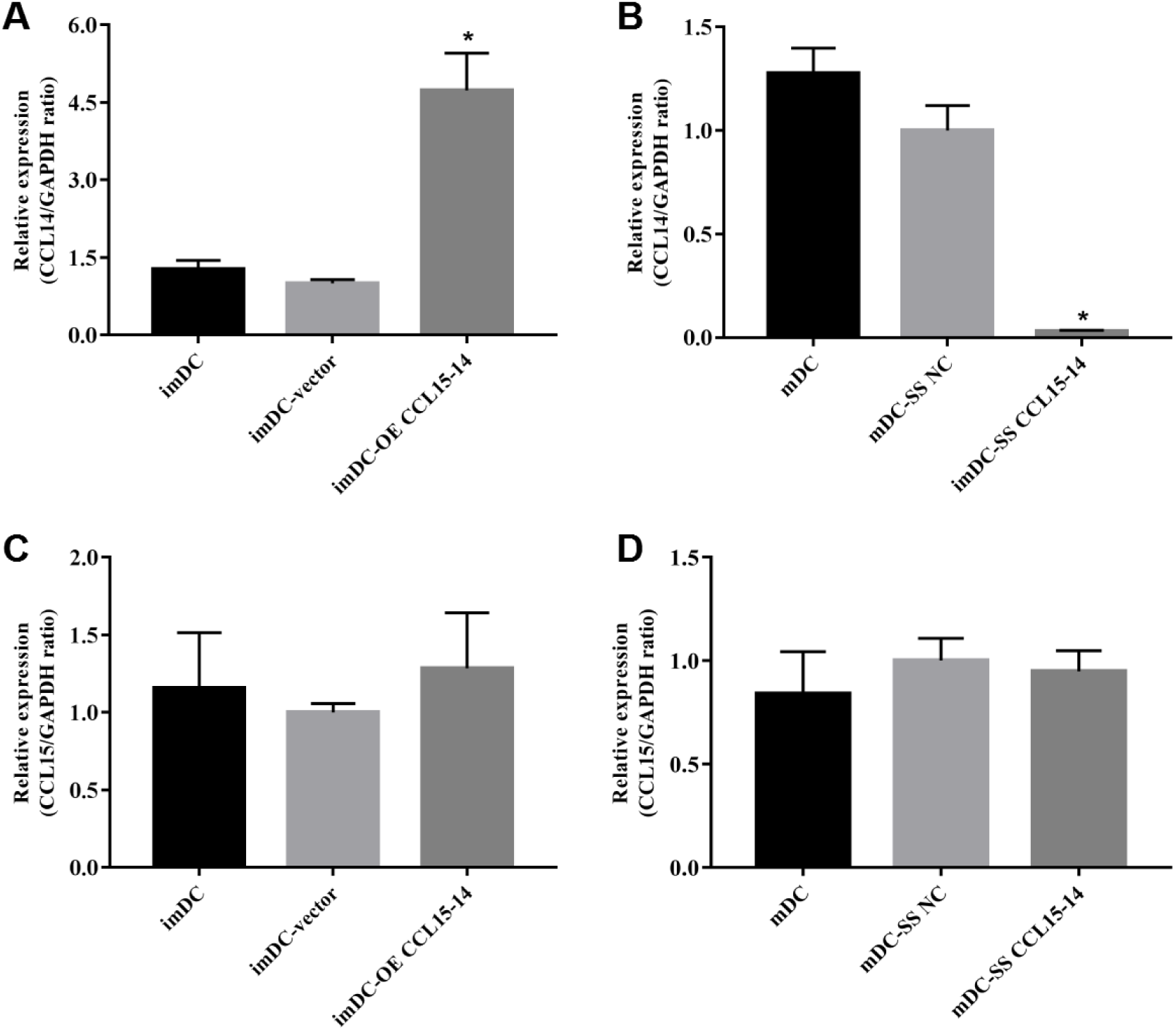
The Expression Levels of CCL14 and CCL15 in moDC were Performed by ELISA Method. Note: n=4 *p<0.05, compared with imDC/mDC group.

**Figure 26.**
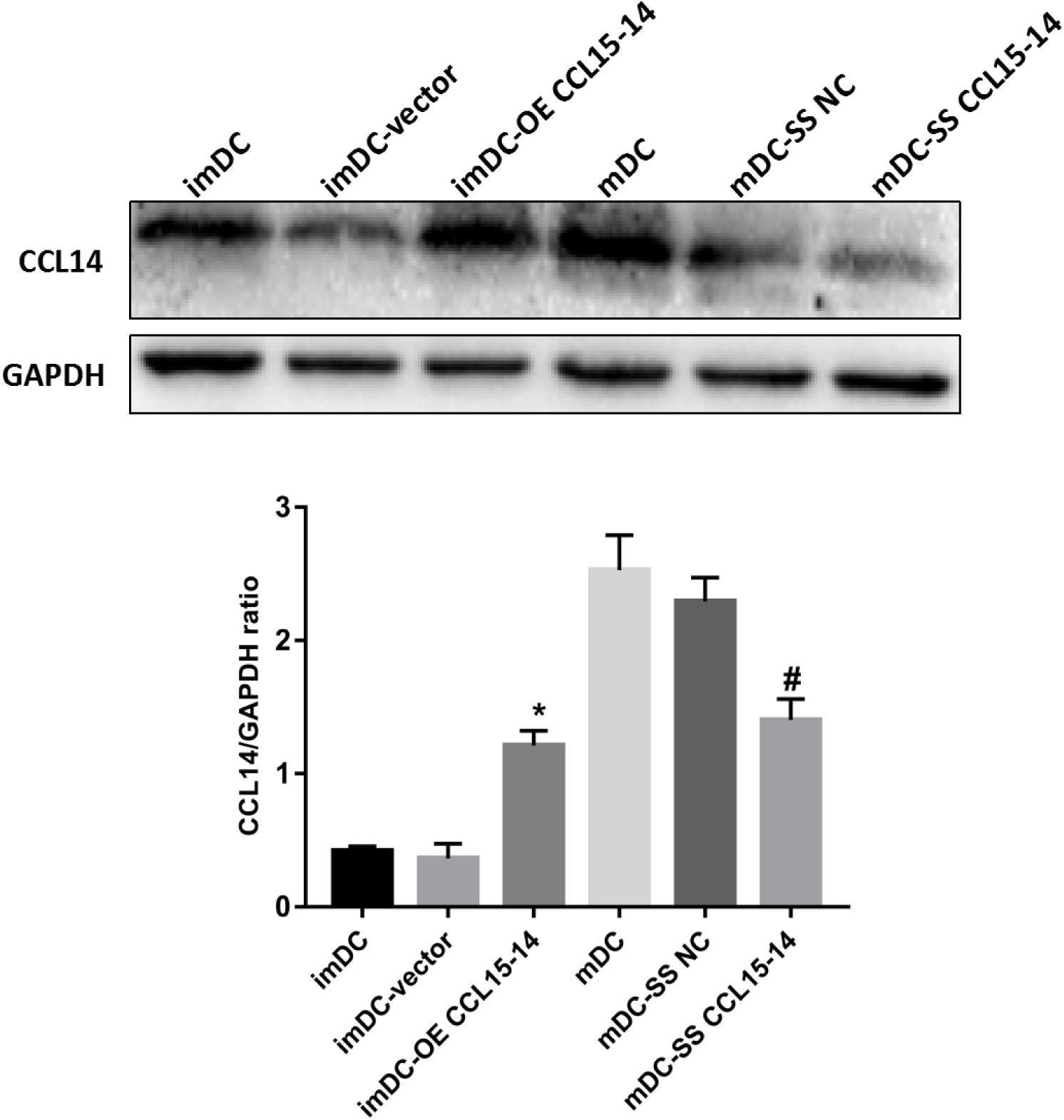
The Expression Level of moDC CCL14 was Performed by Western Blot. Note: n=3 *p<0.05, compared with imDC group, #p<0.05, compared with mDC group.

## Discuss

The present study is the first to explore moDC morphology and function, moDC-derived lncRNA expression differences and the influence of differentially expressed lncRNA on moDC in ACS patients.

In this study, it was found: (1) moDC morphology and function were different in patients with different types of ACS: moDC in patients with acute myocardial infarction had stronger antigen presenting ability, pro-inflammatory secretion function and weaker phagocytosis ability; (2) different types of ACS patients have different expression of moDC derived lncRNA: the number of moDC-derived lncRNA differentially expressed between patients with myocardial infarction (NSTEMI and STEMI) and patients with non-myocardial infarction (UA and CON)was higher. While, there were less lncRNA differentially expressed between CON and UA, and NSTEMI and STEMI. (3) differentially expressed lncRNA (CCL15-CCL14) has a regulatory effect on moDC.

Regarding the relationship between ACS and lncRNA, the first question is that: are there differences in lncRNA expression in patients with ACS? rather than the role of a specific lncRNA in ACS. In 2014, the results of Yang KC et al. showed that 679 and 570 lncRNA were differentially expressed in the ischemic group and the non-ischemic group by collected ventricular tissues for gene sequencing, and lncRNA (rather than mRNA or miRNA) was believed to be involved in heart failure and ventricular remodeling^13^.In 2018, Zhong Z et al. reported that 58 lncRNA were differentially expressed between STEMI and NSTEMI and 106 lncRNA between AMI and non-coronary patients by gene sequencing on circulating monocytes^14,15^. In 2020, the results of Sheng X et al. showed that 552 lncRNA were differentially expressed between AMI and normal subjects by gene sequencing on circulating endothelial cells^16^. The results of present study showed that patients with different types of ACS had different number of lncRNA expressions: the number of differentially expressed lncRNA was higher (CON vs. NST, 49; CON vs ST, 35; UA vs. NST, 115; UA vs ST, 113); However, the number of which between CON and UA (CON vs UA, 3), and between ST and NST (ST vs NST, 4) was lower.

Throughout the above studies, it was found that different types of ACS patients were correlated with the differential expression of lncRNA, and the number of differentially expressed lncRNAs in ACS patients varied greatly among several studies, which was considered to be due to the different sources of lncRNA (monocytes, endothelial cells and cardiomyocytes, etc.).

The second question that needs to be clarified is that: what tissue or cell-derived lncRNA play an important role in the occurrence and progression of ACS?

In present study, moDC was selected as the source of lncRNA for the following four reasons: (1) DC is closely related to the occurrence and development of atherosclerosis and ACS^17^; (2) DC, as an important component of blood cells, plays a role in systemic regulation^18,19^; (3) The number of DC is small, and it is difficult to collect enough DC in ACS patients; (4) Different DC subgroups have significant differences in their functions: at present, DC is divided into four subgroups [20]: A. conventional dendritic cells (cDC), which are mainly used for antigen presentation and activation of T lymphocytes and TH cells to induce cellular and humoral immune effects. B. plasmacytoid dendritic cells (pDC), which produce high-dose type I interferon and thus exert antiviral immunity. C. langerhans cells (LC), which mainly play an immune defense role in the epidermis; D. monocyte-derived dendritic cells (moDC), which play a role in the occurrence and progression of inflammatory diseases^20^. Based on the above, this study selected moDC as the research object to explore the relationship between moDC derived lncRNA and ACS.

The results of this study showed that moDC morphology and function were different in patients with different types of ACS: moDC in patients with acute myocardial infarction had stronger antigen presenting ability, pro-inflammatory secretion function and weaker phagocytosis ability, which was basically consistent with the results of gene sequencing.

NR_027922.3, also known as CCL15-CCL14, is a read-through transcript spanning both CCL14 and CCL15. According to the fold change of gene sequencing (fold change: 4.026332123) and RT-PCR results (among the five candidates lncRNA, NR_027922.3 had the highest expression level), CCL15-CCL14 was selected to study its influence on moDC and the related mechanism. The results showed that lncRNA CCL15-14 could up-regulate or down-regulate the expression levels of DC markers (CD80, CD86, HLA-DR), inflammatory factors (IL-6 and IL-12p70) and target genes (CCL14), and affect the phosphorylation level of PI3K/AKT. As a member of the chemokine family, studies have shown that CCL14 can be up-regulated in dendritic cells to participate in inflammatory responses^21^. CCL14 can also promote the activation of macrophages, promote the phosphorylation level of PI3K/AKT signaling pathway, and positively regulate immune response^22^. Combined with present study, we believe that CCL15-CCL14 activates dendritic cells to participate in the occurrence and development of ACS by regulating the expression level of CCL14 and mediating the phosphorylation of PI3K/AKT.

A surprising result of the study is that: the number of moDC-derived lncRNA differentially expressed between patients with UA and patients with myocardial infarction was the highest, even exceeding the number between CON patients and patients with MI.

The reason may be that patients with UA and patients with MI are both pathological states with many influencing factors, however, CON subject have relatively normal physiological states and fewer influencing factors. From this perspective, it can be inferred that lncRNA differentially expressed between CON and patients with MI is more specific and valuable.

### Limitations

1. Small sample size; (2) The mechanism between CCL15-CCL14 and endothelial cells has not been elucidated.

### Conclusion

There are differences in moDC morphology, function and lncRNA expression in different types of ACS patients, and the differentially expressed lncRNA (CCL15-CCL14) can regulate moDC.

